# Liver regeneration requires reciprocal release of tissue vesicles to govern rapid hepatocyte proliferation

**DOI:** 10.1101/2024.03.03.583167

**Authors:** Si-Qi Ying, Yuan Cao, Ze-Kai Zhou, Xin-Yan Luo, Xiao-Hui Zhang, Ke Shi, Ji-Yu Qiu, Shu-Juan Xing, Yuan-Yuan Li, Kai Zhang, Chen-Xi Zheng, Fang Jin, Yan Jin, Bing-Dong Sui

**Author notes:** **Corresponding authors Prof. Bing-Dong Sui.** Research and Development Center for Tissue Engineering, The Fourth Military Medical University, 145 West Changle Road, Xi’an, Shaanxi 710032, China. **Prof. Yan Jin.** Research and Development Center for Tissue Engineering, The Fourth Military Medical University, 145 West Changle Road, Xi’an, Shaanxi 710032, China. These authors contributed equally to this study (Si-Qi Ying, Yuan Cao).

## Abstract

**Background & Aims:** The liver possesses a remarkable regenerative capacity involving intricate intercellular communication, the mechanisms of which remain poorly understood. Extracellular vesicles (EVs) emerge as important messengers in diverse pathophysiological conditions. However, tissue-derived, cell-specific functional EV populations in regeneration have not been robustly investigated.

**Methods:** Bulk and single-cell RNA sequencing analyses of the regenerating liver after partial hepatectomy (PHx), ultrastructural examinations of *in situ* liver tissue EVs (LT-EVs), and nanoscale and proteomic profiling of hepatocyte-derived tissue EVs (Hep-EVs) were integrated. Targeted inhibition of Hep-EV release *in vivo* was performed *via* AAV-mediated shRNA knockdown of *Rab27a*, and Hep-EVs were intravenously infused for therapeutically use. Gross, histological, and immunofluorescent examinations of livers with evaluating *in vivo* and *ex vivo* hepatocyte proliferation were conducted.

**Results:** LT-EVs contribute to the liver regenerative process after PHx, and hepatocytes serve as the major origin of tissue EVs in the regenerating liver. Moreover, Hep-EVs play an indispensable role to orchestrate liver regeneration, which is strengthened to release with proliferative messages identified after PHx. Mechanistically, Hep-EVs from the regenerating liver reciprocally promote hepatocyte proliferation, which are hallmarked by and function based on the Cyclin dependent kinase 1 (Cdk1) activity. Importantly, replenishment of Hep-EVs from the regenerating liver holds translational promise and rescues insufficient liver regeneration.

**Conclusions:** Our study establishes a functional and mechanistic framework that release of regenerative Hep-EVs govern rapid hepatocyte proliferation through cell cycle control, shedding light on investigation of physiological and endogenous tissue EV populations in organ regeneration and therapy.

## Introduction

The liver is the largest solid organ in the body which critically maintains the metabolism and serves as the central hub for detoxification. In order to cope with daily challenges, the liver has developed a potent, robust, and fine-tuned regenerative capability, which can fully regenerate up-to-2/3 loss of the total parenchyma [1, 2]. The process of liver regeneration involves a complex cellular interplay primarily centered on hepatocytes but also including biliary cells and various non-parenchymal cells (NPCs) [3]. Particularly, in the healthy liver, hepatocytes are mitotically quiescent; whereas following toxic damage or surgical resection, hepatocytes can rapidly enter the cell cycle to proliferate with orchestrating the immune, endothelial, and stromal components to synergistically restore liver mass and function [4–8]. This hepatocyte expansion-related cellular coordination is acknowledged to be mediated by diverse paracrine cytokines, which safeguard efficient liver regeneration as a prerequisite for proper organismal functioning [4, 5, 9]. However, the liver regeneration capacity can be overwhelmed or suppressed under large injury or progressive damages, resulting in hepatic diseases as a global medical challenge [10, 11]. Further deciphering in-depth mechanisms underlying liver regeneration will thus help to devise feasible strategies for hepatic disease therapy.

Extracellular vesicles (EVs) represent an emerging and alternative form of intercellular communication regulating a variety of biological processes, which are functionalized with surface molecules and encapsulated cargoes, such as proteins, nucleic acids, and lipids [12, 13]. Particularly, solid tissues/organs are increasingly recognized as natural reservoirs of endogenous EVs, and tissue-derived EVs play a putative role in reflecting and regulating the tissue health and disease statuses [14]. A recent study has isolated tissue EVs from the kidney and the skin, which have been applied to promote target tissue repair in a donor tissue-specific manner after allogeneic transplantation [15]. It has also been reported that the liver tissue EVs (denoted LT-EVs) isolated from both normal and carbon tetrachloride (CCl_4_)-induced damaged livers accelerate the recovery of liver tissues from CCl_4_-induced hepatic necrosis after administration [16]. However, owing to the heterogeneous nature of tissue resident cells and the imprecision property of the bulk isolation method of tissue EVs, it remains an unmet need to characterize the detailed endogenous EV populations that contribute to tissue physiology and pathology [17–19]. With regard to liver regeneration, EVs derived from the cultured hepatocyte supernatant have been shown to promote liver regeneration after infusion [20]. We have also recently reported that circulatory apoptotic EVs contribute to liver regeneration [21]. Nevertheless, whether LT-EVs from hepatocytes or any other cell types specifically participate in the liver regeneration process has not been evaluated rigorously *in situ*.

Here, we aimed to investigate the role and mechanisms of tissue-derived, cell-specific functional EV population(s) which potentially safeguard liver regeneration. By performing unbiased examinations of the liver transcriptome *via* bulk RNA sequencing (RNA-seq) and the hepatic cell landscape *via* single-cell RNA sequencing (scRNA-seq), we revealed that hepatocytes serve as the key source of intercellular communication in the regenerating liver after partial hepatectomy (PHx), which are programmed to be enriched for biogenesis and release of EVs. Furthermore, we first conducted the ultrastructural observation of LT-EVs *in situ* and established an immunomagnetic sorting protocol analyzing the hepatocyte-specific LT-EVs marked by the featured asialoglycoprotein receptor (ASGPR) (denoted Hep-EVs), which reveal enhanced release at the nanoscale with proliferative information identified at the high-throughput proteomic level during liver regeneration. Then, by targeted inhibition of hepatocyte EV release *in vivo* based on adeno-associated virus (AAV)-mediated short heparin RNA (shRNA) knockdown of the GTPase *Rab27a* expression, we discovered that Hep-EVs are indispensable for orchestrating liver regeneration. Mechanistically, Hep-EVs from the regenerating liver were shown to reciprocally promote proliferation of hepatocytes, which are hallmarked by and function *via* the Cyclin dependent kinase 1 (Cdk1) activity. Importantly, replenishment of Hep-EVs from regenerating livers holds translational promise and rescues insufficient liver regeneration. Therefore, our study establishes a functional and mechanistic framework that release of regenerative Hep-EVs govern rapid hepatocyte proliferation through cell cycle control, shedding light on investigation of physiological and endogenous tissue EV populations in organ regeneration and therapy.

## Materials and Methods

### Animals

C57BL/6 male mice at the 8 weeks of age were used in this study, which were purchased from the Laboratory Animal Center of the Fourth Military Medical University. Mice were group-housed and maintained under specific pathogen-free conditions with standard 12 h light/dark cycles and were kept feeding and drinking *ad libitum*. All animal experiments were performed in compliance with the relevant laws and ethical regulations, following the Guidelines of Intramural Animal Use and Care Committees of the Fourth Military Medical University, approved by the Ethics Committee of the Fourth Military Medical University (No. 2020-003), and following the ARRIVE guidelines.

### PHx surgery

The PHx procedure was performed under a sterile condition [21]. Isoflurane inhalation (R500IP, RWD Life Science, USA) was used to anesthetize the animals. For 2/3 PHx, the median and left lobes of the liver were sequentially resected. The Sham operation was performed by laparotomy. Sampling time points were set at 72 h throughout the study.

### LT-EV collection and Hep-EV isolation

Tissue EVs were collected following our published protocol [22]. Mice were sacrificed after 72 h of Sham or PHx surgeries, and the liver tissue was gently dissociated into tiny pieces before being digestion with the Liberase TM enzyme (LIBTM-RO, Sigma-Aldrich, USA) for 30 min at 37°C, followed by a filtration step through a 0.70-µm pore size filter. The remaining liquid was centrifuged at 300 g for 10 min at 4°C and 2,000 g for 20 min at 4°C to remove cell and tissue debris, and the supernatant was further centrifuged at 16,800 g for 30 min at 4°C to collect LT-EVs. For additional processing, LT-EVs were stained in dark with ASGPR-PE antibody (sc-166633 PE, Santa Cruz Biotechnology, USA; diluted at 1:100) at 4°C for 4 h, and subsequently incubated with the anti-PE magnetic microbeads (130-048-801, Miltenyi Biotec, USA) diluted at 1:16 in MACS buffer (130-091-221, Miltenyi Biotec, USA) at 4°C for 15 min. After passing through the LD column (130-042-901, Miltenyi Biotec, USA), ASGPR^+^ LT-EVs (*i.e.*, Hep-EVs) were captured by the magnet. Hep-EVs were then pushed into fresh sterile tubes and were subjected to two washes with PBS. Hep-EVs were collected after 30 min of centrifugation at 16,800 g at 4°C.

### Hep-EV inhibition and replenishment in vivo

The liver-targeting AAV serotype 8 (AAV8) with a hepatocyte-specific promoter, human α1-antitrypsin promoter (hAATp), and an apolipoprotein E (ApoE) enhancer, was used to specifically deliver shRNA to knockdown *Rab27a* in hepatocytes *in vivo*. Mice were injected with either an AAV vector carrying the *Rab27a* shRNA (pAAV-ApoE/hAATp-*Rab27a*-shRNA, Genechem, China) or a negative control vector (pAAV-ApoE/hAATp-null, Genechem, China) through tail vein at a dose of 1×10^11^ v.g/200 μl per mouse. The AAVs were injected for 4 weeks before PHx and Hep-EV administration. For Hep-EV administration, Hep-EVs in filtered PBS were injected after 28 days of AAV injection *via* the mouse caudal vein at a dose of 5×10^10^ particles/200 μl per mouse for 24 h, followed by PHx surgery.

*Additional materials and methods are described in Supplementary Materials and Methods*.

## Results

### Active involvement of *in situ* LT-EVs in the regeneration process after PHx

To begin, we selected the PHx model which enables the study of liver regeneration without enormous necrosis [1, 2]. As expected, hepatocyte proliferation was confirmed at 72 h after 2/3 PHx by immunofluorescence (IF) staining, with also increased macrophage inflammation and activation of liver sinusoidal endothelial cells (LSECs) and hepatic stellate cells (HSCs), indicating regenerative responses at the organ level (Figure S1A and B). We then performed liver bulk RNA-seq to analyze the potential events occurred during the regenerative process. Within the 9,388 overlapping genes detected between Sham and PHx livers (Figure 1A), 1,009 were differentially expressed genes (DEGs) (Figure 1B, Figure S1C, and Table S1), which demonstrated regenerative characteristics enriched in cell cycle and proliferation regulation, as shown by Gene Oncology (GO) analysis, Kyoto Encyclopedia of Genes and Genomes (KEGG) analysis and Gene Set Enrichment Analysis (GSEA) (Figure S1D-F). Notably, we further discovered remarkable enrichment of DEGs in multiple EV-related GO terms, such as the “Extracellular exosome”, “Extracellular vesicle”, and “Regulation of vesicle-mediated transport” (Figure 1C). Particularly, for the GO terms of “Extracellular exosome” and “Extracellular vesicle”, GSEA revealed significant upregulation in the PHx liver over the Sham control, suggesting EV involvement in the liver regeneration (Figure 1D).

**Figure 1.**
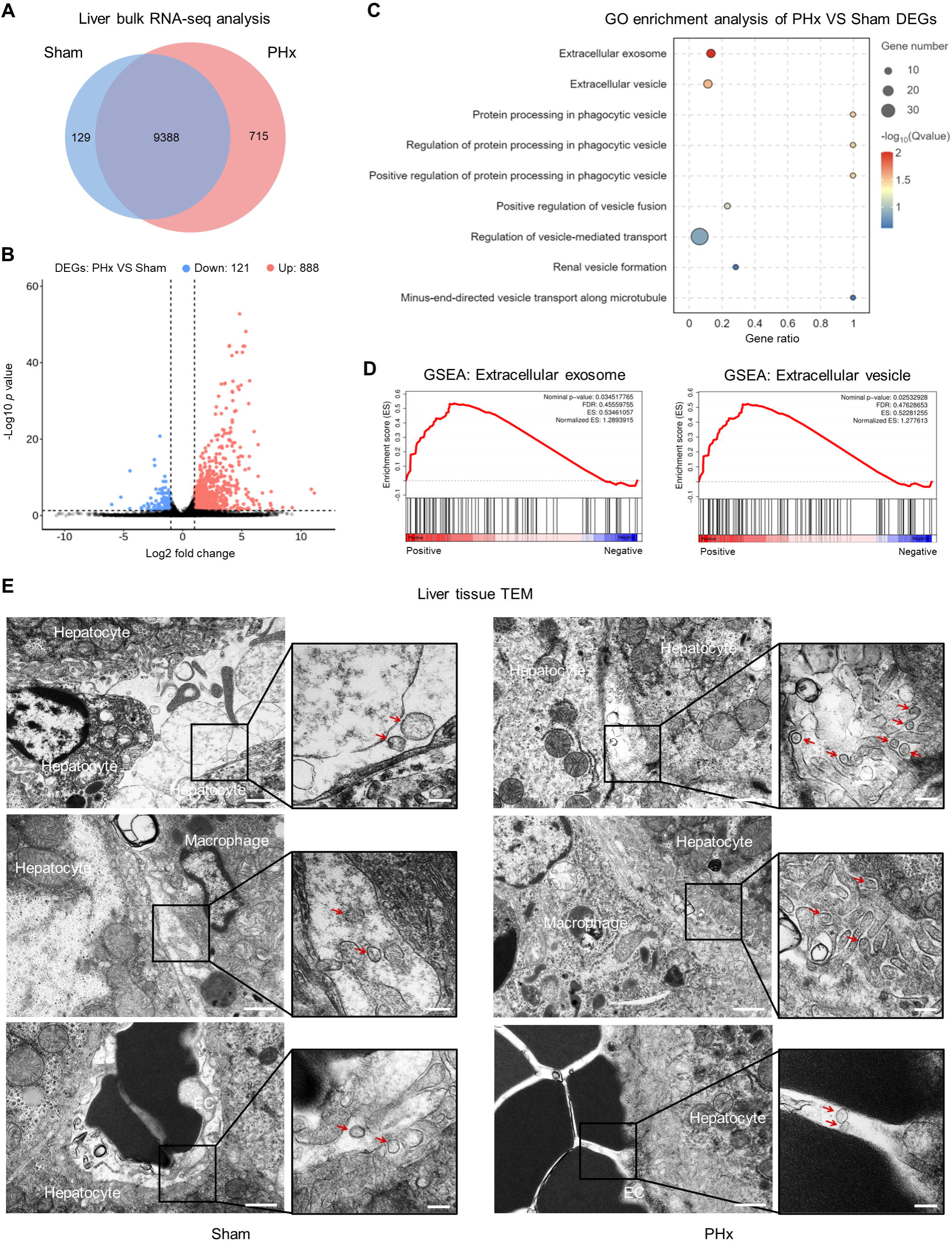
LT-EVs are involved in the liver regenerative process after PHx. **(A)** Venn diagram of transcriptome of PHx and Sham livers. **(B)** Volcano plot of transcriptome of PHx and Sham livers. **(C)** GO terms of DEGs related to EVs enriched in PHx over Sham livers. **(D)** GSEA analysis of DEGs between PHx and Sham livers for the GO terms “extracellular exosome” and “extracellular vesicle”. **(E)** TEM analysis of Sham and PHx liver tissues. Hepatocytes, macrophages, and ECs were identified with featured morphologies. Red arrows indicating LT-EVs in the extracellular interstitial space. Bars: 1 µm (low magnification) and 200 nm (high magnification).

Next, we intended to validate that whether LT-EVs indeed participated in regeneration. Although not proved rigorously, *in situ* EVs putatively exist in the tissue extracellular space and reflect pathophysiological features of the tissue [14, 15]. However, current studies examine tissue EVs mainly *via* enzymatic digestion and isolation methods, resulting in loss of spatiotemporal information and potential compromise of EV amount and integrity [17–19]. Existing literatures are also limited on the evaluation of *in situ* EVs in tissue regeneration. To detect the *bona fide* EVs in the liver, we conducted transmission electron microscopy (TEM) analysis of liver tissues. In tissue sections of PHx, compared to those of Sham, hepatocytes were enlarged and the interstitial space among neighbored hepatocytes was narrowed, in accordance with the hypertrophic phenotype in regeneration (Figure 1E) [23]. Intriguingly, in the extracellular space adjacent to hepatocytes, vesicle-like nanoparticles were observed, which were presence in the Sham liver and demonstrated a dramatic increase in abundance in the PHx liver (Figure 1E). In tissue extracellular spaces between hepatocytes and macrophages or endothelial cells (ECs), nanosized vesicles were also detected in Sham and PHx livers (Figure 1E). Together, these findings first indicate the active involvement of *in situ* EVs in liver regeneration.

### Hepatocytes are the major origin of LT-EVs during liver regeneration

Next, we examined the origin of LT-EVs in the liver regenerative process. Accordingly, we re-analyzed two scRNA-seq data of the mouse liver respectively from Sham and PHx conditions (GSM4572241 and GSM4572244). Cell clustering using t-distributed stochastic neighbor embedding (tSNE) plot visualization identified 8 distinct liver cell types, in which the proportion of hepatocytes decreased after PHx (Figure 2A and B). Despite decreased proportion, pseudotime analysis showed that hepatocytes from Sham and PHx livers fell into distinct stage branches with a few transitioning cells, indicating a cellular state shift (Figure S2A). A total of 3,524 DEGs between Sham and PHx hepatocytes were identified (Figure 2C), which were categorized based on the functional annotations of GO and KEGG databases, such as the GO term “cellular anatomical entity” and the KEGG term “Transport and catabolism” (Figure S2B and C). In-depth functional implications of the DEGs further demonstrated remarkable enrichment of EV-related GO terms, particularly associated with EV assembly and secretion (Figure 2D). These results suggest that hepatocytes might be programmed for biogenesis and release of EVs in liver regeneration.

**Figure 2.**
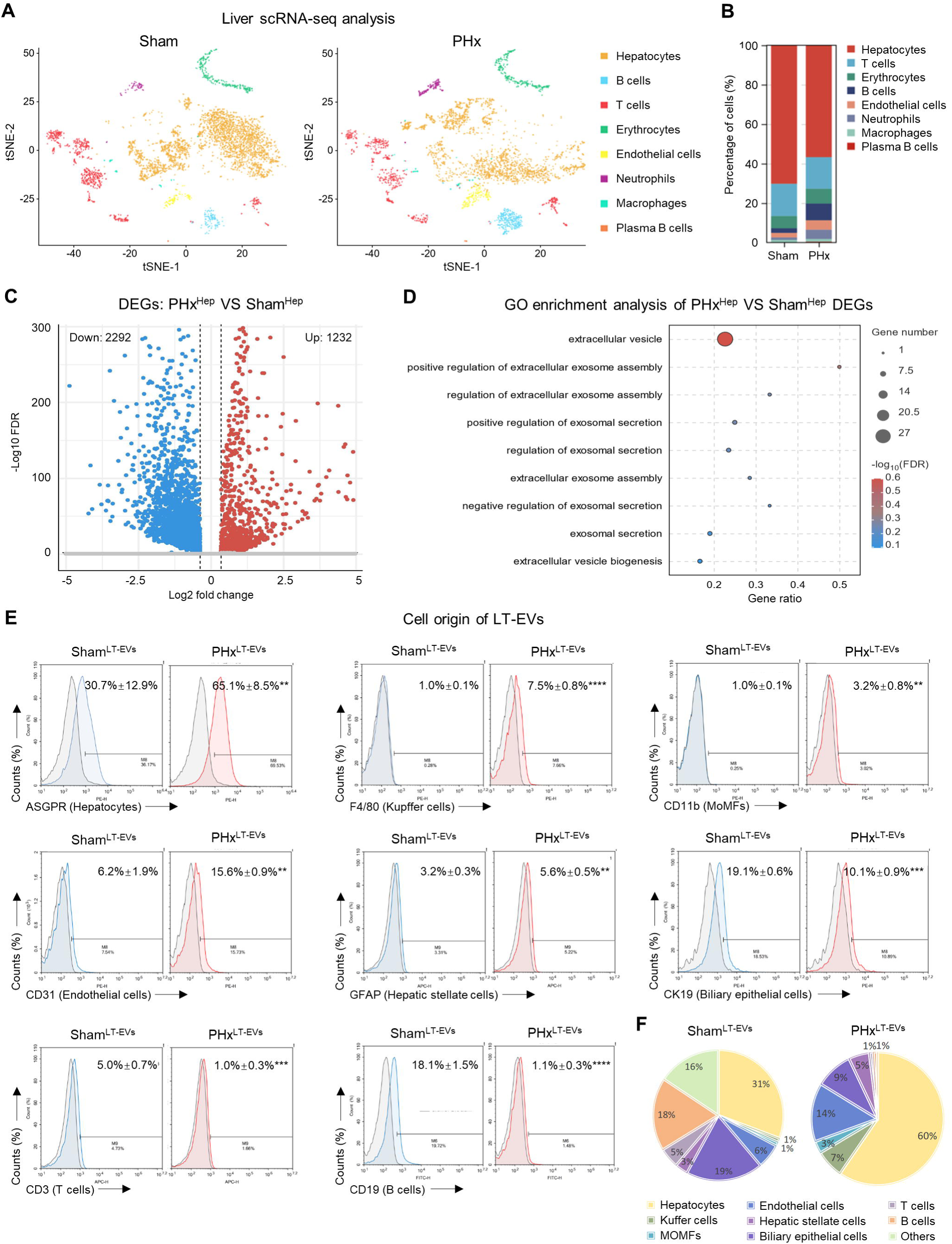
Hepatocytes are the major origin of LT-EVs during liver regeneration. **(A)** tSNE plots displaying the distribution of different cell types in Sham and PHx livers. Source data were derived from the GSM4572241 and GSM4572243 series in the GEO database. **(B)** Cell type proportion in Sham and PHx livers. **(C)** Volcano plot showing DEGs between PHx and Sham hepatocytes. **(D)** GO enrichment analysis of DEGs in PHx over Sham hepatocytes. **(E)** Flow cytometric analysis of Sham and PHx LT-EVs with surface exposure of ASGPR, F4/80, CD11b, CD31, GFAP, CK19, CD3, and CD19. Corresponding isotype control groups were used to distinguish positively stained EVs. Mean ± SD. n = 3 per group. **, *p* < 0.01; ***, *p* < 0.001; ****, *p* < 0.0001; two-tailed Student’s unpaired *t* tests. **(F)** Pie charts illustrating the quantification of LT-EV constitution originated from different cell types using mean values of (E) normalized to percentages.

We then performed flow cytometric analysis of collected LT-EVs to investigate the surface antigens inherited from their parental cells. Interestingly, ASGPR-marked LT-EVs derived from hepatocytes were the largest ingredients of total LT-EVs at the physiological state (a mean percentage at about 31%), which demonstrated an approximately two-fold increase after PHx (Figure 2E). Other main components of Sham LT-EVs included those from biliary epithelial cells (marked by cytokeratin 19, CK19) and B cells (marked by CD19), which, nevertheless, were decreased after PHx (Figure 2E). Another LT-EV subpopulations with increased proportions during the liver regeneration process were from F4/80-marked Kupffer cells and CD31-marked ECs, while LT-EVs from liver-infiltrated monocyte-derived macrophages (MoMFs, marked by CD11b), HSCs (marked by glial fibrillary acidic protein, GFAP), and T cells (CD3) were only around or below 5% percentages of total constitution (Figure 2E). Correspondingly, we mapped the previously unrecognized cell origin of LT-EVs (Figure 2F). Taken together, these results suggest that hepatocytes are the major origin of LT-EVs and serve as the key source of intercellular EV communication in the regenerating liver.

### Hep-EV release is indispensable for orchestrating liver regeneration

The above findings prompted us to investigate that whether Hep-EVs were functionally required for liver regeneration. To address this question, we constructed and intravenously (*i.v.*) administered the pAAV-ApoE/hAATp-*Rab27a*-shRNA to specifically inhibit hepatocyte release of EVs *in vivo* (Figure 3A). As expected, compared to the PHx group and PHx mice injected with the control vector, AAV8-mediated sh*Rab27a* delivery successfully reduced LT-EV amount during liver regeneration (Figure 3B). To specifically analyze Hep-EVs, we established a general protocol for collecting and characterizing antigen-specific tissue EVs by integrating our previous methods of immunomagneto-activated sorting of the marked EV population from cultured cells and isolation of traceable EVs from the tissue (Figure S3A) [22, 24]. Using this method with adopting the ASGPR surface marker, we then validated the suppressed release of Hep-EVs *in vivo* after pAAV-ApoE/hAATp-*Rab27a*-shRNA delivery (Figure 3B). Importantly, targeted inhibition of Hep-EV release substantially reduced the survival rate of mice after the PHx surgery, with only 25% of survival at 72 h, indicating diminished liver regeneration (Figure 3C). Furthermore, gross analysis of livers from even survived mice at 72 h, with histological examination and quantification of liver weight over body weight (LW/BW) ratio, confirmed retarded regeneration without tissue alterations after AAV8-mediated *Rab27a* knockdown (Figure 3D and E). Indeed, IF staining further showed that pAAV-ApoE/hAATp-*Rab27a*-shRNA delivery prevented hepatocytes from entering into proliferation after the PHx challenge, suggesting the functional importance of Hep-EVs for liver parenchymal recovery (Figure 3F and G). Besides, AAV8-mediated *Rab27a* inhibition led to suppressed inflammation of liver macrophages with reduced normal and activated LSECs, as well as decreased quiescent HSCs yet increased activated HSCs, indicating the necessity of Hep-EVs coordinating immune, endothelial, and mesenchymal responses in liver regeneration (Figure 3F and G). Taken together, these findings highlight that Hep-EV release is indispensable for orchestrating liver regeneration.

**Figure 3.**
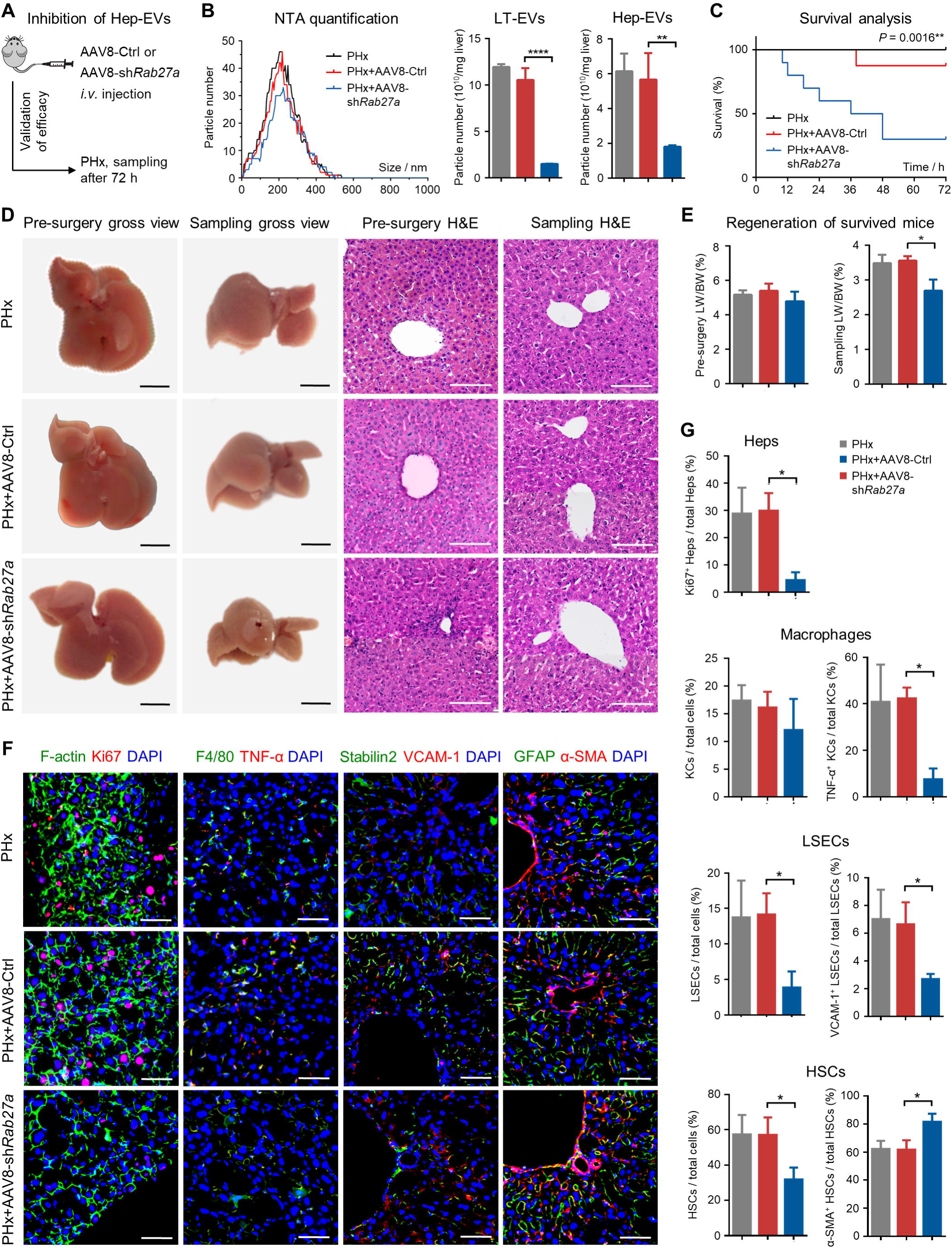
Hep-EV release is indispensable for orchestrating liver regeneration. **(A)** Schematic diagram of the study design using AAV8-shRNA to inhibit hepatocyte *Rab27a via i.v.* injection prior to PHx. **(B)** Quantification of particle number of LT-EVs and Hep-EVs determined by NTA. Hep-EVs were immunomagetically sorted from LT-EVs. Mean ± SD. n = 3 per group. **, *p* < 0.01; ****, *p* < 0.0001; one-way ANOVA with Turkey’s post-hoc tests. **(C)** Kaplan-Meier survival analysis of mice. n = 8 per group. Log-rank test was used. **(D)** The gross view images and H&E staining of liver tissues. Bars = 500 mm (gross view) and 100 mm (H&E). **(E)** Quantification of liver weight over body weight ratio. Mean ± SD. n = 3 per group. *, *p* < 0.05; one-way ANOVA with Turkey’s post-hoc tests. **(F)** IF staining of hepatocyte proliferation, macrophage inflammation, LSEC and HSC activation in the liver. Bars = 50 μm. **(G)** Quantification of percentages of proliferative hepatocytes, total and inflammatory macrophages, total and activated LSECs and HSCs in the liver. Mean ± SD. n = 3 per group. *, *p* < 0.05; one-way ANOVA with Turkey’s post-hoc tests.

### Hep-EVs reveal enhanced release with proliferative information in liver regeneration

Next, we investigated the characteristics of Hep-EVs in liver regeneration. As expected, flow cytometric data confirmed that the ASGPR^+^ percentage of LT-EVs was 33.79% before immunomagnetic sorting, while the percentage decreased to 0.10% in ASGPR^-^ LT-EVs and increased to 98.34% in sorted Hep-EVs (Figure 4A and Figure S3B). After validation of the efficacy, we adopted this approach to investigate LT-EVs and Hep-EVs derived from Sham and PHx mice, with the quality of EVs evaluated according to the Minimal information for studies of extracellular vesicles 2023 (MISEV2023) [25]. Nanoparticle tracking analysis (NTA) demonstrated that the diameters of LT-EVs from both Sham and PHx mice ranged from 50 to 500 nm and peaked at 150-200 nm (Figure 4B). Notably, compared to Sham mice, PHx mice produced significantly more amount of LT-EVs, as quantified by calculating the particle number by NTA and the protein content by the BCA assay (Figure 4C). Further observation by TEM showed that LT-EVs exhibited the typical cup-shape morphology with membranous structure in Sham and PHx (Figure 4D). Furthermore, after immunomagnetic sorting, Hep-EVs from Sham and PHx mice still showed characteristic particle distribution in NTA (Figure 4E), and Hep-EVs derived from the PHx liver revealed higher yield than the Sham counterparts (Figure 4F). Taken together, these results suggest that hepatocytes strengthen EV release in liver regeneration.

**Figure 4.**
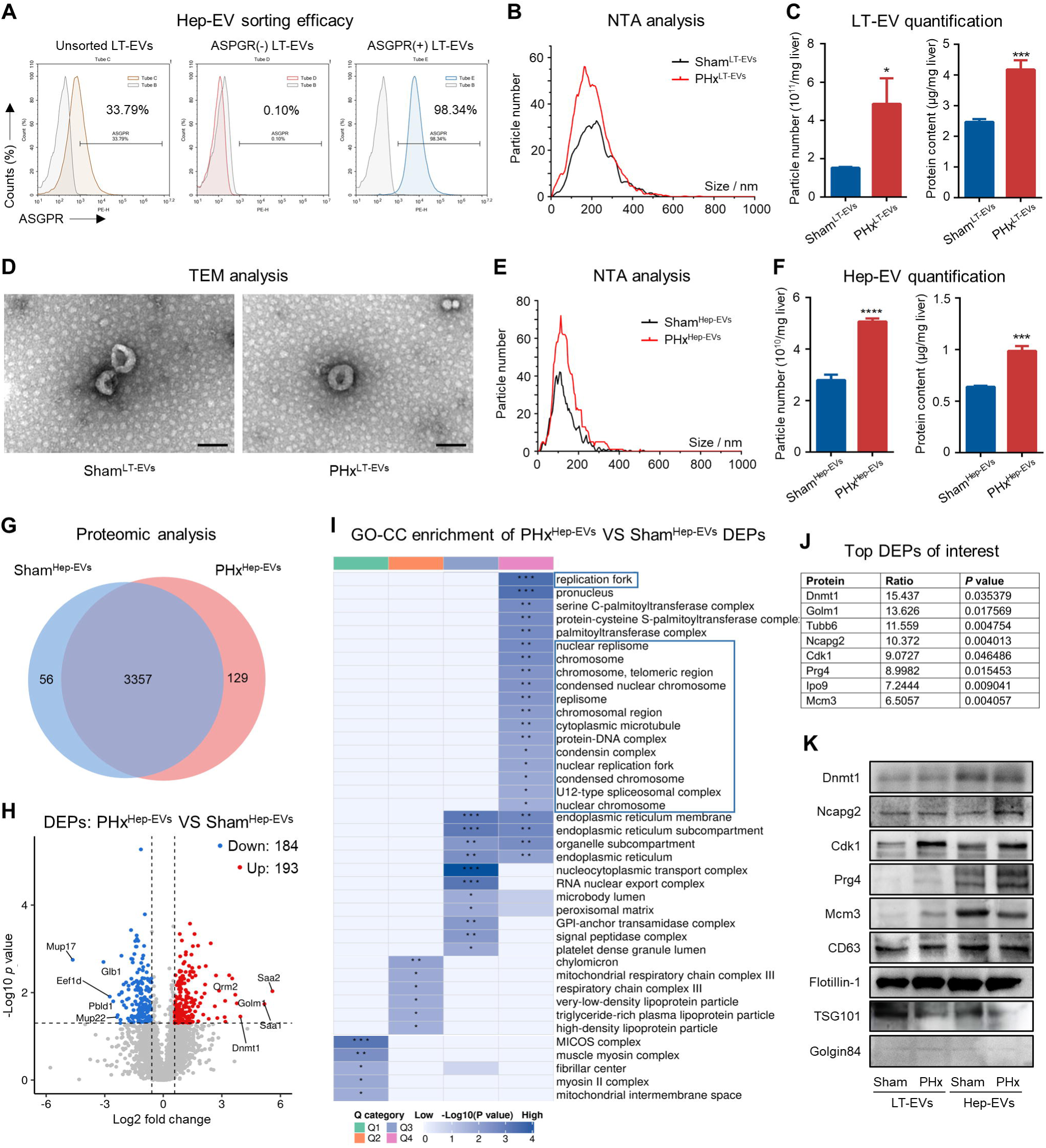
Hep-EVs reveals enhanced release with proliferative information during liver regeneration. **(A)** Flow cytometric analysis showing ASGPR surface expression on LT-EVs before and after immunomagnetic sorting. **(B)** Size distribution of LT-EVs analyzed by NTA. **(C)** Quantification of particle number of LT-EVs by NTA and protein amount of LT-EVs determined by the BCA assay. Mean ± SD. n = 3 per group. *, *p* < 0.05; ***, *p* < 0.001; one-way ANOVA with Turkey’s post-hoc tests. **(D)** Representative TEM image of LT-EVs from Sham and PHx livers. Bars = 250 nm. **(E)** Size distribution of sorted Hep-EVs analyzed by NTA. **(F)** Quantification of particle number of sorted Hep-EVs by NTA and protein amount determined by the BCA assay. Mean ± SD. n = 3 per group. ***, *p* < 0.001; ****, *p* < 0.0001; one-way ANOVA with Turkey’s post-hoc tests. **(G)** Venn diagram of proteome of PHx and Sham Hep-EVs. **(H)** Volcano plot of proteome of PHx and Sham Hep-EVs. **(I)** GO terms in the Cellular Component category of DEPs enriched in PHx over Sham Hep-EVs. **(J)** Western blot analysis of protein marker expression of LT-EVs and Hep-EVs. **(K)** Top-ranked DEPs of interest in Hep-EVs and Western blot validation of expression.

We then examined the protein cargo profiles of Hep-EVs from Sham and PHx livers by performing the liquid chromatography with tandem mass spectrometry (LC-MS/MS). A total of over 3000 proteins were identified (Table S2). The Venn diagram illustrated the presence of 3357 shared proteins between Sham and PHx Hep-EVs, with respective 56 and 129 distinct proteins in each group (Figure 4G). Among the shared proteins, 387 proteins were identified as the differentially expressed proteins (DEPs; fold change > 1.5 and *p* value < 0.05), and 193 DEPs were upregulated in PHx Hep-EVs with 184 DEPs downregulated (Figure 4H), showing clear separation and distinct protein signatures between Sham and PHx Hep-EVs (Figure S4A). GO enrichment analysis of DEPs in the Cellular Component (GO-CC) category further demonstrated that annotations related to the proliferative process, such as “replication fork”, “chromosomal, telomeric region”, and “cytoplasmic microtubule”, were particular regulated terms in Hep-EVs during liver regeneration (Figure 4I). Regarding the functionality of DEPs, they were also notably associated with “DNA replication” and “Cell cycle” in the KEGG enrichment analysis, as well as certain mitosis-related terms in the Biological Process (GO-BP) and Molecular Function (GO-MF) categories of GO databases (Figure S4B-D).

The above findings identified liver regeneration-associated cargo profiles of Hep-EVs centered on proliferative regulation. To validate these data, we selected several top DEPs of interest for Western blot analysis (Figure 4J). Not surprisingly, both LT-EVs and Hep-EVs from Sham and PHx mice expressed representative EV markers, such as the tetraspanin surface molecule CD63, the lipid raft protein Flotillin-1, and the endosomal sorting complex required for transport (ESCRT)-associated protein tumor suppressor gene 101 (TSG101) (Figure 4K). Meanwhile, the Golgi component protein Golgin84, a negative EV marker, was not detected in either LT-EVs or Hep-EVs (Figure 4K). Interestingly, PHx-derived Hep-EVs were verified to highly express the matrix protein Proteoglycan 4 (Prg4) and the Non-SMC condensin II complex subunit G2 (Ncapg2), which mediates the microtubule-attachment process to accelerate mitosis [26], despite not being enriched with the epigenetic enzyme DNA methyltransferase 1 (Dnmt1) and the DNA helicase Minichromosome maintenance complex component 3 (Mcm3) [27] (Figure 4K). It was notable that Cdk1, the critical kinase to promote the G2/M and G1/S transitions [28], was especially enriched in both LT-EVs and Hep-EVs during liver regeneration (Figure 4K). Taken together, these findings highlight that Hep-EVs reveal enhanced release with proliferative information in liver regeneration.

### Hep-EVs from the regenerating liver reciprocally promote hepatocyte proliferation ***via* Cdk1 activity**

We kept focusing on these DEPs and intended to dissect the potential relationship between Hep-EV proteomic alterations and transcriptomic changes of their parental cells. By integrating the scRNA-seq data and the proteomic data, we performed a nine-quadrant analysis to distinguish cellular and vesicular expression patterns under liver regeneration. The graph depicted the same expression patterns in a particular interested quadrant of P3, showing upregulation in both cases (Figure 5A). GO enrichment analysis of this quadrant revealed association of terms with cellular activity, especially the terms “epidermal growth factor receptor binding” and “cyclin B1-CDK1 complex”, suggesting proliferation-related functionality (Figure 5B). Intriguingly, data suggested that organelle-related proteins were uniquely upregulated in Hep-EVs irrespective of the cellular transcriptome (Figure S5A), whereas metabolism-related terms showed consistent downregulation in hepatocytes and their EVs during liver regeneration (Figure S5B). Together, these results indicate that EVs diversify hepatocyte-derived information meanwhile inherit potential proliferative regulation capability during liver regeneration.

**Figure 5.**
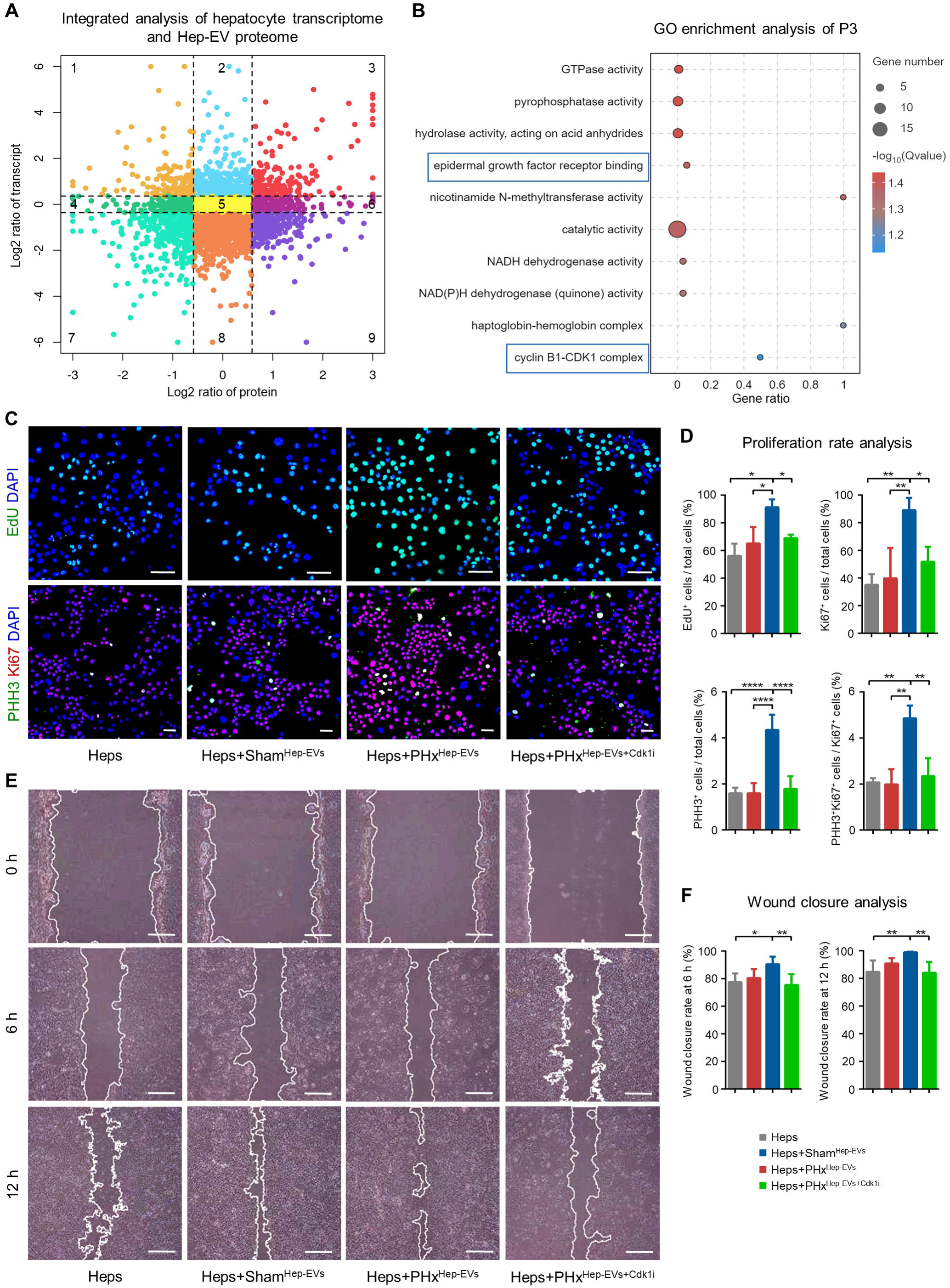
Hep-EVs from the regenerating liver reciprocally promote proliferation of hepatocytes *via* Cdk1 activity. **(A)** Nine-quadrant diagram showing integrated analysis of hepatocyte transcriptome from scRNA-seq and Hep-EV proteome. **(B)** GO enrichment analysis of the P3 quadrant showing consistent upregulation of hepatocyte genes and Hep-EV proteins. **(C)** EdU assay and IF staining of PHH3 with Ki67 for proliferative hepatocytes treated with Hep-EVs and the Cdk1 inhibitor RO-3306. Bars = 100 µm. **(D)** Quantification of percentages of EdU^+^, Ki67^+^, PHH3^+^ hepatocytes and PHH3^+^Ki67^+^ hepatocytes among Ki67^+^ hepatocytes. n = 3 per group. *, *p* < 0.05; **, *p* < 0.01; ***, *p* < 0.001; ****, *p* < 0.0001; one-way ANOVA with Turkey’s post-hoc tests. **(E)** Representative images of wound closure assay of hepatocytes treated with Hep-EVs and the Cdk1 inhibitor RO-3306. Bars = 250 µm. **(F)** Quantification of wound closure rates of hepatocytes. n = 3 per group. *, *p* < 0.05; **, *p* < 0.01; one-way ANOVA with Turkey’s post-hoc tests.

We then deciphered that whether PHx-derived Hep-EVs were indeed able to regulate cell proliferation for liver regenerative repair. Although that many liver cell populations were affected by Hep-EV inhibition *in vivo* (Figure 3F), we focused on hepatocytes as putative reciprocal targets of Hep-EVs given the critical role they plays to regenerate the liver based on effective proliferation control [1]. Furthermore, considering the proliferative information carried by PHx-derived Hep-EVs, particularly the Cdk1 enrichment (Figure 4K and 5B), we tested that whether PHx-derived Hep-EVs regulated target cell mitosis mediated by Cdk1 activity. To this end, cultured hepatocytes were treated with Hep-EVs from Sham and PHx livers, and PHx-derived Hep-EVs were preconditioned with or without a chemical inhibitor of Cdk1. Expectedly, 5-ethynyl-2’-deoxyuridine (EdU) labeling analysis and Ki67 IF staining demonstrated that PHx-derived Hep-EVs significantly promoted DNA synthesis of cultured hepatocytes, while Sham-derived Hep-EVs failed to do so (Figure 5C and D). IF staining of Ki67 with Phosphohistone H3 (PHH3), which marks cells in late G2 and M phases [29], further revealed the effect of PHx-derived Hep-EVs, but not their Sham counterparts, to accelerate hepatocyte mitosis (Figure 5C and D). The effects of PHx-derived Hep-EVs on hepatocyte proliferation were suppressed by the Cdk1 inhibitor pretreatment, as shown by reduced EdU, Ki67, and PHH3 positive cell percentages (Figure 5C and D), which was particularly related to Cdk1 inhibition given the Cdk1 function to facilitate the G1/S and G2/M transitions [28]. The cellular wound closure assay confirmed the beneficial effects of PHx-derived Hep-EVs, rather than Sham-derived Hep-EVs, on the healing capability of hepatocytes, which were also based on Cdk1 activity (Figure 5E and F). Collectively, these findings suggest that Hep-EVs from the regenerating liver reciprocally promote hepatocyte proliferation *via* Cdk1 activity.

### Replenishment of Hep-EVs from the regenerating liver rescues insufficient liver regeneration

Finally, we investigated that whether Hep-EVs were of translational promise to promote liver regeneration upon infusion. As a first proof-of-concept experiment, we intravenously administered liver-isolated Hep-EVs to replenish diminished *in vivo* Hep-EVs in recipient AAV8-treated *Rab27a*-knockdown mice (Figure 6A). Importantly, survival analysis showed that Hep-EV replenishment was remarkably effective to promote liver regeneration after the PHx surgery, and mice injected with PHx-derived Hep-EVs exhibited higher survival rate than those receiving Sham-derived EVs (Figure 6B). Further gross view with histological analysis of livers from survived mice of each group confirmed that PHx-derived Hep-EVs significantly promoted liver regeneration upon infusion (Figure 6C and D). The effects of Hep-EVs to promote hepatocyte proliferation after replenishment were also discovered by IF staining, in which infusion of both Sham-derived and PHx-derived Hep-EVs increased the Ki67-positive hepatocyte percentages *in vivo*, especially PHx-derived Hep-EVs (Figure 6C and D). Taken together, these results suggest that replenishment of Hep-EVs from the regenerating liver is able to rescue insufficient liver regeneration.

**Figure 6.**
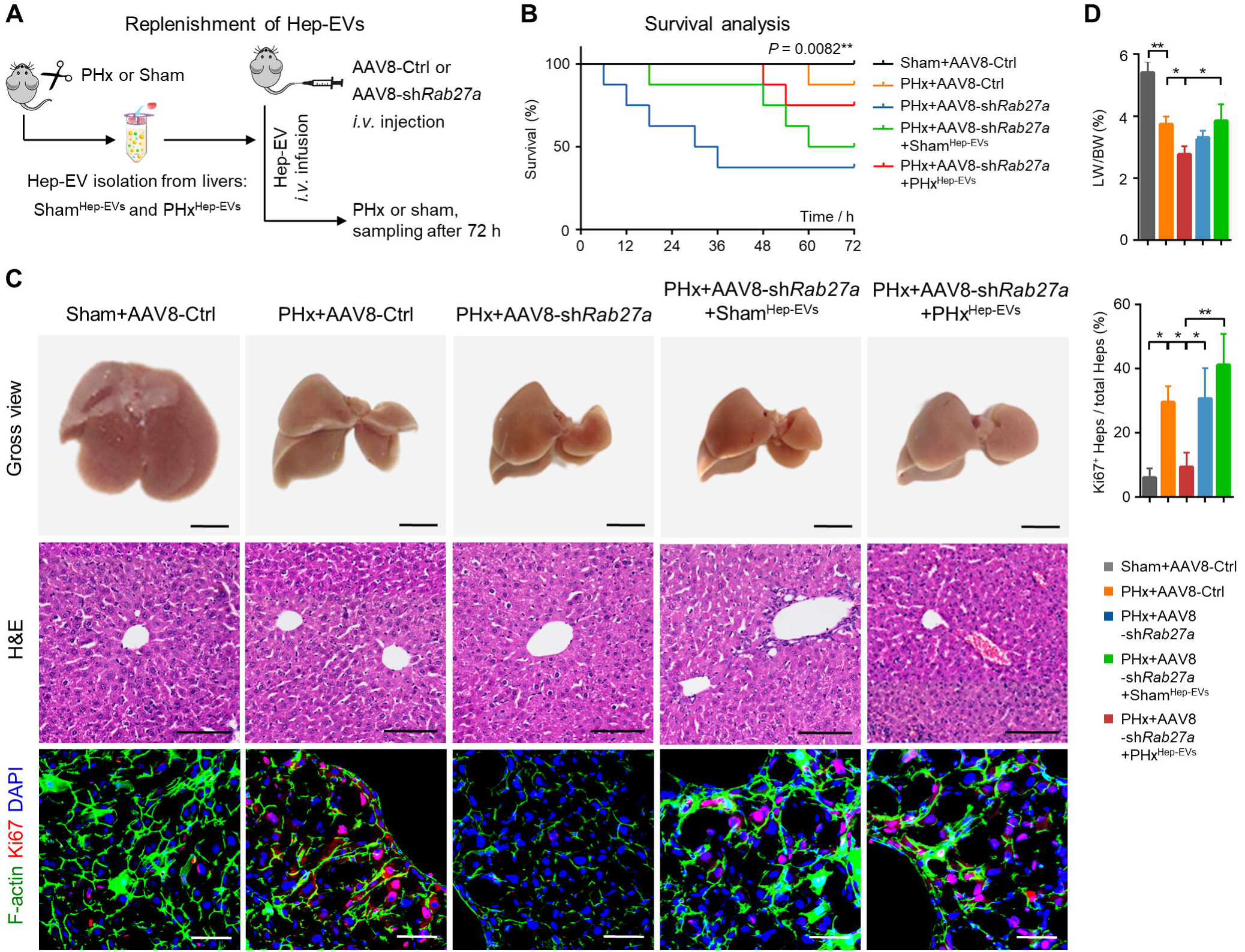
Replenishment of Hep-EVs from the regenerating liver rescues insufficient liver regeneration. **(A)** Schematic diagram of the study design using Hep-EV infusion as a replenishment into AAV8-sh*Rab27a*-treated mice before liver regeneration analysis. **(B)** Kaplan-Meier survival analysis of mice. n = 8 per group. Log-rank test was used. **(C)** Gross view liver images, H&E staining of liver tissues, and IF staining of hepatocyte proliferation. Bars = 500 mm (gross view), 100 mm (H&E), and 50 μm (IF). **(D)** Quantification of liver weight over body weight ratio and percentages of proliferative hepatocytes. Mean ± SD. n = 3 per group. *, *p* < 0.05; **, *p* < 0.01; one-way ANOVA with Turkey’s post-hoc tests.

## Discussion

The liver is responsible for maintaining metabolism and detoxification of the body and is highly regenerative [1, 2]. The process of liver regeneration involves complex intercellular communication [3]. However, the mechanism of this elaborate functional interaction has not been thoroughly understood. EVs are biologically active paracrine messengers which act dynamically in various pathophysiological states [12, 13]. In the present study, we mapped the transcriptomic landscape of liver regeneration at both the bulk and the single cell levels, which highlighted the crucial role of hepatocyte-centered intercellular communication based on EVs. Our investigation further uncovered that liver regeneration promoted the release of LT-EVs, particularly Hep-EVs, which were confirmed both *in situ* and *ex vivo* with inheriting regulated regenerative cues. Notably, Hep-EVs were demonstrated to promote hepatocyte proliferation in a reciprocal mode based on the Cdk1 activity, which played a necessary role *in vivo* to safeguard liver regeneration in the physiology and upon therapeutic infusion. For the first time, this study provides an integrated phenotypic and functional characterization of the hepatocyte-specific tissue EVs in liver regeneration, offering insights into the biological and mechanistic aspects of the regenerating liver.

The liver possesses the remarkable ability to fully regenerate after undergoing partial resection [2]. Studies have documented that liver regeneration involves a combination of subsequent hepatocyte division, hypertrophy, and proliferation [8, 23]. Our sequencing data further suggested significant influences of liver regeneration on the metabolic processes of hepatocytes, which are largely suppressed in the regenerating liver. Indeed, the metabolic reprogramming after PHx represent an important event, and the transcriptome sequencing combined with metabolomic analysis has indicated detailed metabolic changes during liver regeneration [30, 31]. Our findings, together with others’ findings and despite preliminary, suggest that hepatocytes might sacrifice metabolic activity for safeguarding regenerative expansion, and that hepatocytes are equipped with flexible metabolic machinery able to adapt dynamically to tissue regeneration [31]. It is also noteworthy that our data revealed involvements of the immune, endothelial and stromal cell components in liver regeneration, which are in accordance with the current understanding [3, 10, 32], and that communication between hepatocytes and various cell populations through EVs potentially contributed to liver regeneration. Future research is needed to examine in-depth the multimodal paracrine regulation mechanisms *via* Hep-EVs that safeguard liver regeneration.

Researchers worldwide have recently dedicated significant efforts to investigating EVs. These membranous nanoparticles are discovered to escape from the autophagy-lysosomal pathway and be release into the circulation and the extracellular space of tissues [33, 34]. Particularly, studies have shown the potential of tissue EVs in promoting regeneration, and LT-EVs have been employed to expedite the hepatic disease recovery process [15, 16, 35]. However, the specific cell origin of tissue EVs responsible for the action remains unknown. In this study, we established an immunomagnetic assay to specifically isolate Hep-EVs in the regenerating liver. As phenotypically assessed, the isolation procedure did not exert significant impact on EV morphology and size distribution, suggesting that the properties of these tissue-derived EVs were largely maintained. With advantages of potent specificity and efficiency [36, 37], utilization of this approach in the EV research will benefit biological and mechanistic investigations of endogenous tissue EVs, as well as for diagnostic and therapeutic applications. In addition, this approach can be applied to gain profound insights into crucial subsets of EVs not only derived from specific cell sources but also possessing featured membrane proteins. Future examinations should elucidate distinct functionalities of LT-EVs from diverse cell sources in liver health and disease.

The pivotal role EVs play in regulating liver health and disease has been suggested, but the function of endogenous LT-EVs remains not fully understood [38]. In a recent work, it has been documented that Interferon regulatory factor 1 (IRF1)-Rab27a-regulated EVs promote liver ischemia/reperfusion injury through surface oxidized phospholipids (OxPL) activation of neutrophils [35]. In our study, we have also discovered that the regenerative Hep-EVs are released based on Rab27a control, while it remains to be investigated the upstream molecular regulator of Rab27a following the PHx challenge. For the EV diversity contributing to liver regeneration, we have previously uncovered that circulatory apoptotic vesicles (apoVs) safeguard liver regeneration through facilitating the organelle assembly of hepatocytes [21]. Others have unraveled that PHx-induced apoVs stimulate neutrophils to secrete regenerative growth factors [39]. Here, we first reveal that Hep-EVs are functionally required to govern rapid hepatocyte proliferation *in vivo* and *in vitro*, which further show translational promise to promote liver regeneration upon systemic infusion. It has recently been reported that the EV cargoes are not randomly but specifically selected [40]. Notably, the Cdk1-based cell cycle control mechanism will have particular implications as potential pharmacological targets [28]. Therefore, the effects of Hep-EV infusion on promoting liver regeneration are valuable for the translational use of tissue EVs in therapies.

In summary, our findings provide the first knowledge that hepatocyte-specific tissue vesicles are phenotypically involved in and functionally required for liver regeneration. Our study paves an avenue for in-depth biological and mechanistic research of regenerative Hep-EVs and sheds light on physiological and endogenous tissue EV populations in organ regeneration and therapy.

## Acknowledgements

This work was supported by grants from the National Natural Science Foundation of China (82371020 to B.D.S., 82301028 to C.X.Z., 81930025 to Y.J. and 82170988 to F.J.) and the Young Science and Technology Rising Star Project of Shaanxi Province (2023KJXX-027 to B.D.S.). We are grateful for the assistance of the National Experimental Teaching Demonstration Center for Basic Medicine (AMFU). We thank PTM Biolabs Co., Ltd (Hangzhou, China) for help in analyzing the proteomic data. We thank Mr. Tao Liu (Gene Denovo Biotechnology Co, Ltd, 790 Guangzhou, China) for his assistance in analyzing the single-cell sequencing data.

## Author contributions

S.Q.Y. and Y.C. contributed equally to the design and conduction of the study and drafted the manuscript. Z.K.Z. and X.Y.L. analyzed and interpreted data. X.H.Z. contributed to the transcriptomic and scRNA-seq experiments. K.S. performed partial hepatectomy. J.Y.Q. contributed to NTA and WB experiments. S.J.X. contributed to H&E and IF experiments.

Y.Y.L. contributed to TEM and flow cytometry analysis. K.Z. contributed to the proteomic experiment. C.X.Z. and F.J. contributed to the discussion and interpretation of data. Y.J. and B.D.S. conceived the project, designed and supervised the experiments. All authors revised the manuscript and approved the final version.

## Data and materials availability

All data included in this study are available upon reasonable request of corresponding authors.

---

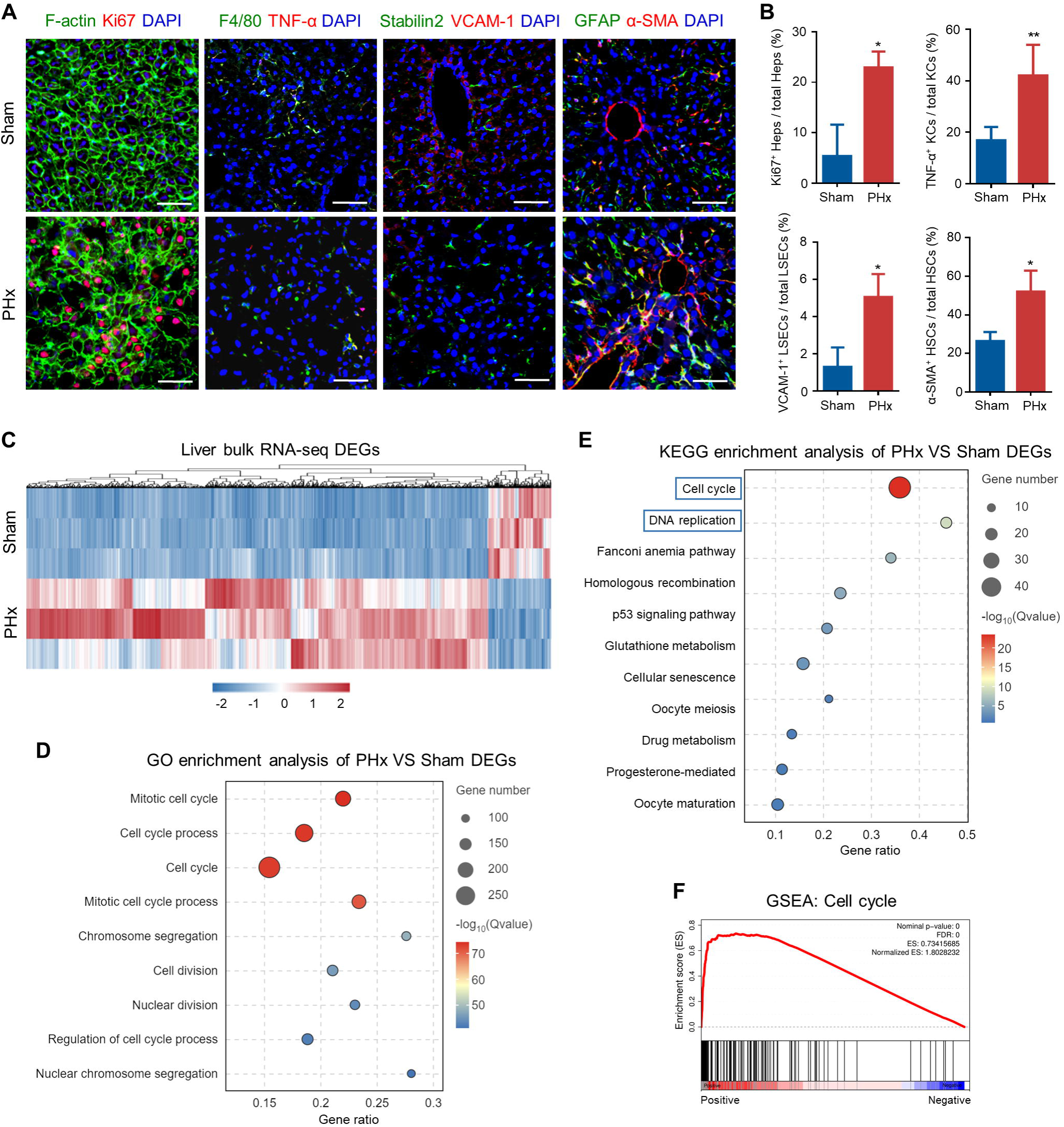

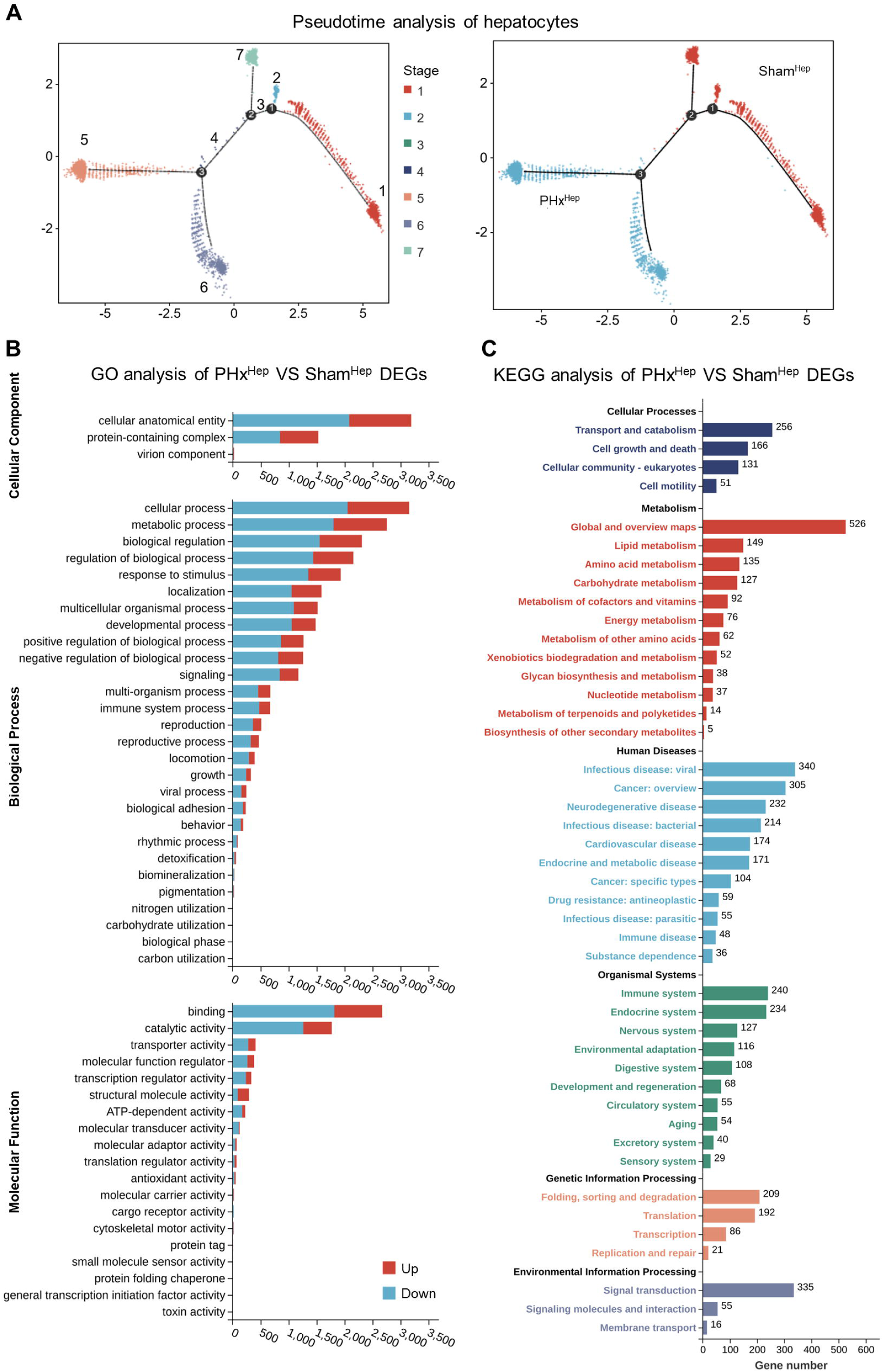

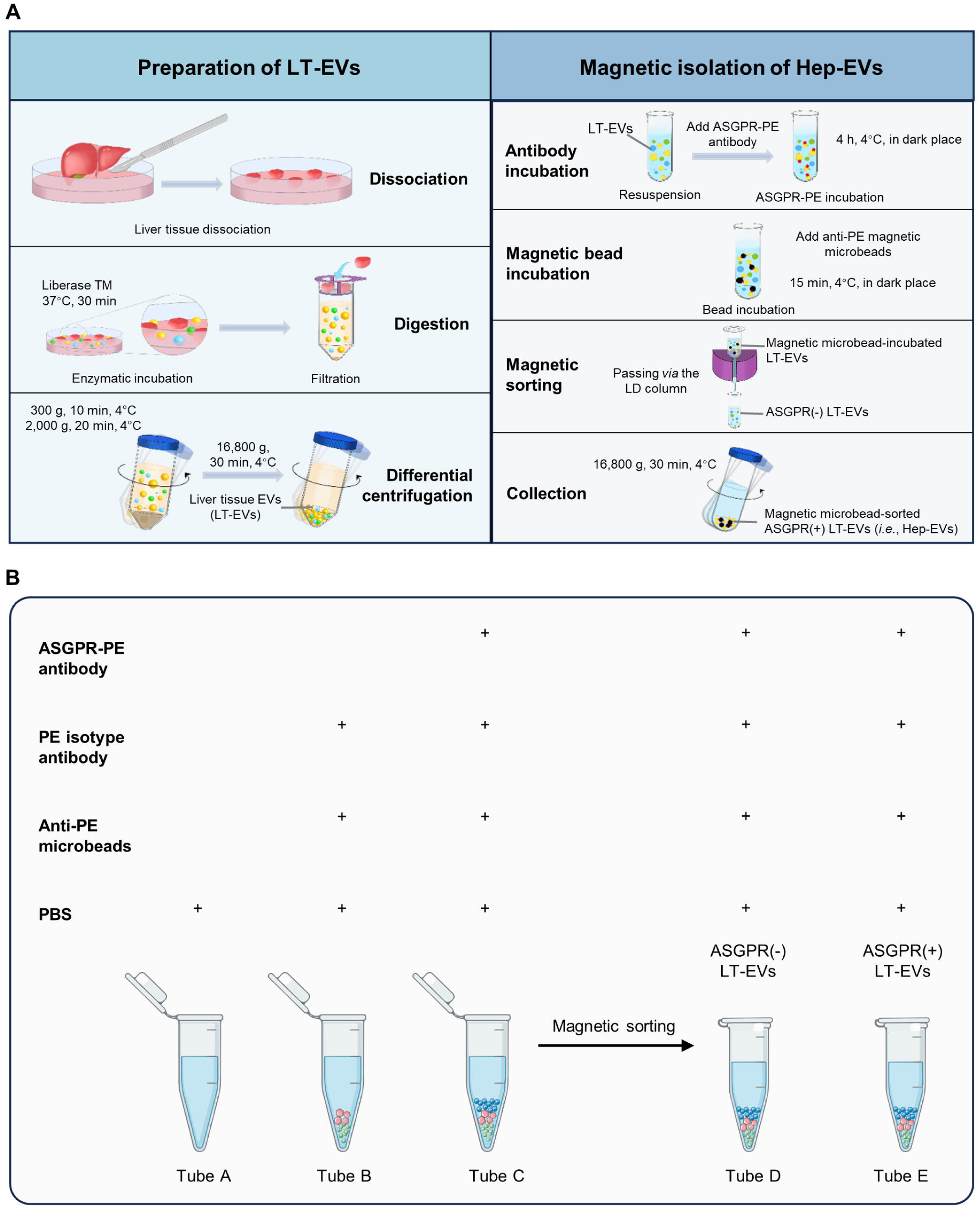

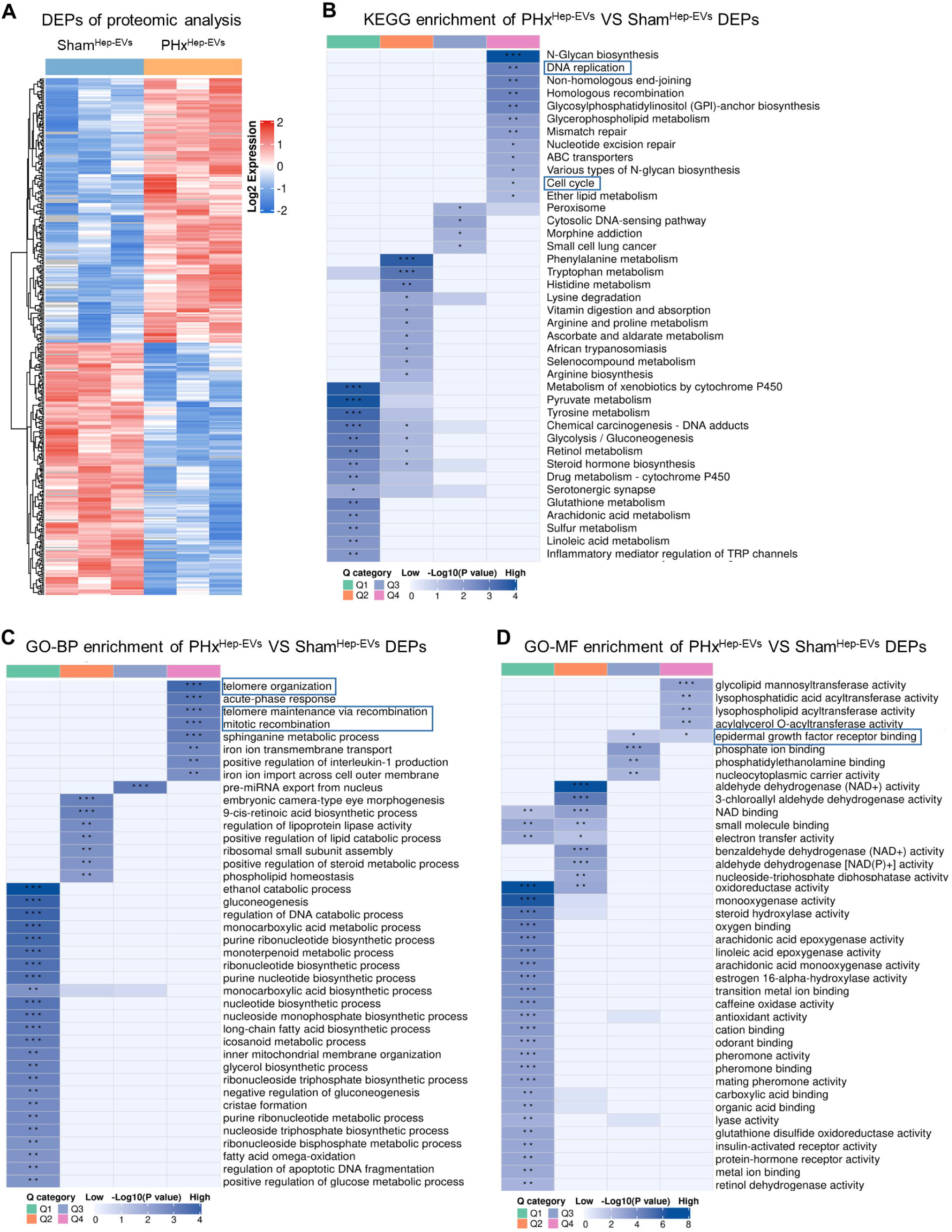

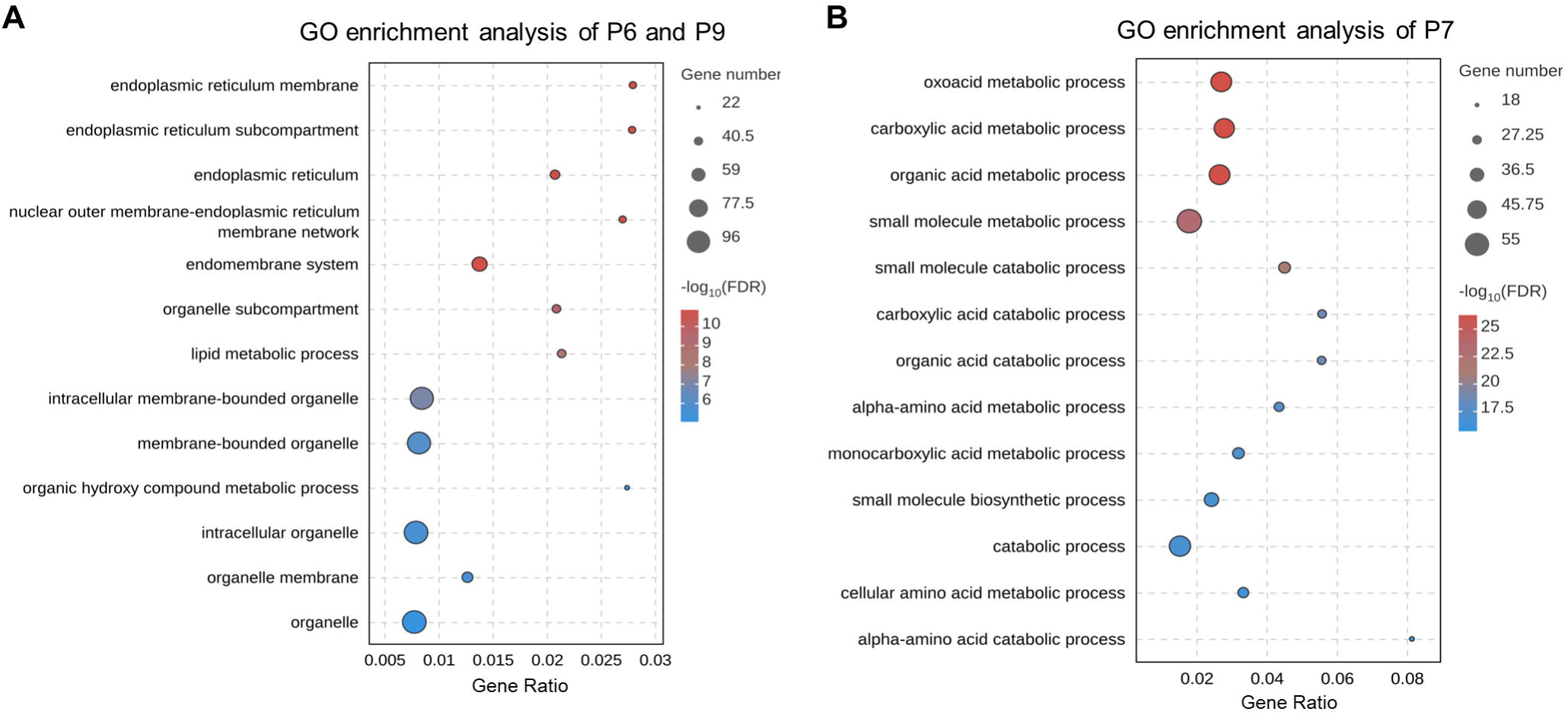

## References

[1] Michalopoulos GK, Bhushan B. Liver regeneration: biological and pathological mechanisms and implications. Nat Rev Gastroenterol Hepatol 2021;18:40–55.

[2] Michalopoulos GK. Liver regeneration after partial hepatectomy: critical analysis of mechanistic dilemmas. Am J Pathol 2010;176:2–13.

[3] Campana L, Esser H, Huch M, Forbes S. Liver regeneration and inflammation: from fundamental science to clinical applications. Nat Rev Mol Cell Biol 2021;22:608–624.

[4] Ait Ahmed Y, Fu Y, Rodrigues RM, He Y, Guan Y, Guillot A, et al. Kupffer cell restoration after partial hepatectomy is mainly driven by local cell proliferation in IL-6-dependent autocrine and paracrine ma nners. Cell Mol Immunol 2021;18:2165–2176.

[5] Shimizu H, Miyazaki M, Wakabayashi Y, Mitsuhashi N, Kato A, Ito H, et al. Vascular endothelial growth factor secreted by replicating hepatocytes induces sinusoidal endothelial cell proliferation during regeneration after partial hepatectomy in rats. Journal of hepatology 2001;34:683–689.

[6] Xiao T, Meng W, Jin Z, Wang J, Deng J, Wen J, et al. miR-182-5p promotes hepatocyte-stellate cell crosstalk to facilitate liver regeneration. Communications biology 2022;5:771.

[7] Song J, Ma J, Liu X, Huang Z, Li L, Li L, et al. The MRN complex maintains the biliary-derived hepatocytes in liver regeneration through ATR-Chk1 pathway. NPJ Regenerative medicine 2023;8:20.

[8] Yanger K, Knigin D, Zong Y, Maggs L, Gu G, Akiyama H, et al. Adult hepatocytes are generated by self-duplication rather than stem cell differentiation. Cell stem cell 2014;15:340–349.

[9] Mei Y, Thevananther S. Endothelial nitric oxide synthase is a key mediator of hepatocyte proliferation in response to partial hepatectomy in mice. Hepatology 2011;54:1777–1789.

[10] Forbes SJ, Newsome PN. Liver regeneration - mechanisms and models to clinical application. Nat Rev Gastroenterol Hepatol 2016;13:473–485.

[11] Stravitz RT, Lee WM. Acute liver failure. Lancet 2019;394:869–881.

[12] S ELA, Mager I, Breakefield XO, Wood MJ. Extracellular vesicles: biology and emerging therapeutic opportunities. Nature reviews Drug discovery 2013;12:347–357.

[13] Yanez-Mo M, Siljander PR, Andreu Z, Zavec AB, Borras FE, Buzas EI, et al. Biological properties of extracellular vesicles and their physiological functions. J Extracell Vesicles 2015;4:27066.

[14] Qin B, Hu XM, Su ZH, Zeng XB, Ma HY, Xiong K. Tissue-derived extracellular vesicles: Research progress from isolation to application. Pathol Res Pract 2021;226:153604.

[15] Lou P, Liu S, Wang Y, Lv K, Zhou X, Li L, et al. Neonatal-Tissue-Derived Extracellular Vesicle Therapy (NEXT): A Potent Strategy for Precision Regenerative Medicine. Adv Mater 2023;35:e2300602.

[16] Lee J, Kim SR, Lee C, Jun YI, Bae S, Yoon YJ, et al. Extracellular vesicles from in vivo liver tissue accelerate recovery of liver necrosis induced by carbon tetrachloride. J Extracell Vesicles 2021;10:e12133.

[17] Verweij FJ, Balaj L, Boulanger CM, Carter DRF, Compeer EB, D’Angelo G, et al. The power of imaging to understand extracellular vesicle biology in vivo. Nature methods 2021;18:1013–1026.

[18] Crescitelli R, Lasser C, Lotvall J. Isolation and characterization of extracellular vesicle subpopulations from tissues. Nat Protoc 2021;16:1548–1580.

[19] Vella LJ, Scicluna BJ, Cheng L, Bawden EG, Masters CL, Ang CS, et al. A rigorous method to enrich for exosomes from brain tissue. J Extracell Vesicles 2017;6:1348885.

[20] Nojima H, Freeman CM, Schuster RM, Japtok L, Kleuser B, Edwards MJ, et al. Hepatocyte exosomes mediate liver repair and regeneration via sphingosine-1-phosphate. Journal of hepatology 2016;64:60–68.

[21] Sui B, Wang R, Chen C, Kou X, Wu D, Fu Y, et al. Apoptotic Vesicular Metabolism Contributes to Organelle Assembly and Safeguards Liver Homeostasis and Regeneration. Gastroenterology 2024.

[22] Cao Y, Qiu JY, Chen D, Li CY, Xing SJ, Zheng CX, et al. Isolation and Analysis of Traceable and Functionalized Extracellular Vesicles from the Plasma and Solid Tissues. J Vis Exp 2022.

[23] Miyaoka Y, Ebato K, Kato H, Arakawa S, Shimizu S, Miyajima A. Hypertrophy and unconventional cell division of hepatocytes underlie liver regeneration. Curr Biol 2012;22:1166–1175.

[24] Zheng C, Sui B, Zhang X, Hu J, Chen J, Liu J, et al. Apoptotic vesicles restore liver macrophage homeostasis to counteract type 2 diabetes. J Extracell Vesicles 2021;10:e12109.

[25] Welsh JA, Goberdhan DCI, O’Driscoll L, Buzas EI, Blenkiron C, Bussolati B, et al. Minimal information for studies of extracellular vesicles (MISEV2023): From basic to advanced approaches. J Extracell Vesicles 2024;13:e12404.

[26] Wang Q, Li Z, Zhou S, Li Z, Huang X, He Y, et al. NCAPG2 could be an immunological and prognostic biomarker: From pan-cancer analysis to pancreatic cancer validation. Front Immunol 2023;14:1097403.

[27] Alvarez S, Diaz M, Flach J, Rodriguez-Acebes S, Lopez-Contreras AJ, Martinez D, et al. Replication stress caused by low MCM expression limits fetal erythropoiesis and hematopoietic stem cell functionality. Nature communications 2015;6:8548.

[28] Lukasik P, Zaluski M, Gutowska I. Cyclin-Dependent Kinases (CDK) and Their Role in Diseases Development-Review. Int J Mol Sci 2021;22.

[29] Elmaci I, Altinoz MA, Sari R, Bolukbasi FH. Phosphorylated Histone H3 (PHH3) as a Novel Cell Proliferation Marker and Prognosticator for Meningeal Tumors: A Short Review. Appl Immunohistochem Mol Morphol 2018;26:627–631.

[30] Li Z, Peng B, Chen S, Li J, Hu K, Liao L, et al. Transcriptome sequencing and metabolome analysis reveal the metabolic reprogramming of partial hepatectomy and extended hepatectomy. BMC genomics 2023;24:532.

[31] Caldez MJ, Van Hul N, Koh HWL, Teo XQ, Fan JJ, Tan PY, et al. Metabolic Remodeling during Liver Regeneration. Developmental cell 2018;47:425–438 e425.

[32] Matchett KP, Wilson-Kanamori JR, Portman JR, Kapourani CA, Fercoq F, May S, et al. Multimodal decoding of human liver regeneration. Nature 2024;630:158–165.

[33] Cao Y, Ying SQ, Qiu XY, Guo J, Chen C, Li SJ, et al. Proteomic analysis identifies Stomatin as a biological marker for psychological stress. Neurobiol Stress 2023;22:100513.

[34] Perrin P, Janssen L, Janssen H, van den Broek B, Voortman LM, van Elsland D, et al. Retrofusion of intralumenal MVB membranes parallels viral infection and coexists with exosome release. Curr Biol 2021;31:3884–3893 e3884.

[35] Yang MQ, Du Q, Goswami J, Varley PR, Chen B, Wang RH, et al. Interferon regulatory factor 1-Rab27a regulated extracellular vesicles promote liver ischemia/reperfusion injury. Hepatology 2018;67:1056–1070.

[36] Tang P, Thongrom B, Arora S, Haag R. Polyglycerol-Based Biomedical Matrix for Immunomagnetic Circulating Tumor Cell Isolation and Their Expansion into Tumor Spheroids for Drug Screening. Adv Healthc Mater 2023;12:e2300842.

[37] Shin HS, Park J, Lee SY, Yun HG, Kim B, Kim J, et al. Integrative Magneto-Microfluidic Separation of Immune Cells Facilitates Clinical Functional Assays. Small 2023:e2302809.

[38] Balaphas A, Meyer J, Sadoul R, Morel P, Gonelle-Gispert C, Buhler LH. Extracellular vesicles: Future diagnostic and therapeutic tools for liver disease and regeneration. Liver Int 2019;39:1801–1817.

[39] Brandel V, Schimek V, Gober S, Hammond T, Brunnthaler L, Schrottmaier WC, et al. Hepatectomy-induced apoptotic extracellular vesicles stimulate neutrophils to secrete regenerative growth factors. Journal of hepatology 2022;77:1619–1630.

[40] Dixson AC, Dawson TR, Di Vizio D, Weaver AM. Context-specific regulation of extracellular vesicle biogenesis and cargo selection. Nat Rev Mol Cell Biol 2023;24:454–476.

