## supplementary information for "Liver regeneration requires reciprocal release of tissue vesicles to govern rapid hepatocyte proliferation"

**^*^Correspondence to:**

**This file includes:**

Supplementary Materials and Methods

Figures S1 to S5, with their legends

Captions of Tables S1 and S2

**Supplementary Materials and Methods**

*Liver bulk RNA-seq analysis*

The total RNA of the whole liver was extracted using the Trizol reagent kit (Invitrogen, USA) following the manufacturer's protocol. RNA quality was assessed on an Agilent 2100 Bioanalyzer (Agilent Technologies, USA) and checked using RNase free agarose gel electrophoresis. After total RNA was extracted, eukaryotic mRNA was enriched by Oligo (dT) beads. Then the enriched mRNA was fragmented into short fragments using fragmentation buffer and reversely transcribed into cDNA using NEBNext Ultra RNA Library Prep Kit for Illumina (NEB #7530, New England Biolabs, USA). The purified double-stranded cDNA fragments were end repaired, a base added, and ligated to Illumina sequencing adapters. The ligation reaction was purified with the AMPure XP Beads (1.0X) and amplified by polymerase chain reaction (PCR). The resulting cDNA library was sequenced using Illumina Novaseq6000 by Gene Denovo Biotechnology Co. (Guangzhou, China). The genes/transcripts with the parameter of false discovery rate (FDR) below 0.05 and absolute fold change ≥ 2 were considered differentially expressed genes/transcripts. Further functional analysis was conducted based on the GO and KEGG databases or through GSEA enrichment analysis.

*scRNA-seq microarray dataset*

scRNA-seq microarray datasets were screened and downloaded from the Gene Expression Omnibus (GEO). The GSM4572241 and GSM4572243 series were obtained from the GEO database for analysis of DEGs. Further functional analysis was conducted based on the GO and KEGG databases.

*Cell clustering*

The cell-by-gene matrices for each sample were individually imported to Seurat version 3.1.1 for downstream analysis. The Seurat was used to perform expression quality check, data normalization, t-distributed stochastic neighbor embedding (tSNE) plot analysis, cell clustering, cluster visualization, and cell type annotation.

*Pseudotemporal trajectory analysis*

Single-cell pseudotemporal trajectory was analyzed using matrices of cells and gene expressions by Monocle (Version 2.10.1). The space was reduced with two dimensions and cells were ordered (sigma = 0.001, lambda = NULL, param.gamma = 10, tol = 0.001). The trajectory was visualized using a tree-like structure, including tips and branches.

*Proteomic analysis*

Protein lysates of Hep-EVs in Sham and PHx groups were prepared and subjected to liquid chromatography with tandem mass spectrometry (LC-MS/MS) analysis on an Orbitrap Exploris™ 480 mass spectrometer with a NanoSpray III ion source. The raw data were analyzed using the Proteome Discoverer system (v2.4.1.15). Proteins were identified by comparing with the Uniport database with a false discovery rate set at 0.01 for both peptides and proteins. Proteins were quantified using the default parameters in MaxQuant. Among the identified proteins, 516 proteins were DEPs (fold change > 1.5 and p-value < 0.05). Proteins were included for further functional analysis based on GO and KEGG databases.

*Cell culture and assays*

The murine AML12 hepatocytes (ATCC CRL-2254) were cultured in Dulbecco’s modified Eagle’s medium with F-12 (DMEM F-12, Gibco, USA) at 37°C with 5% CO_2_. The medium was supplemented with 10% fetal bovine serum (FBS; HyClone, USA), 1% ITS liquid medium supplement (Sigma Aldrich, USA), 1% 100 μg/ml of penicillin and streptomycin (HyClone, USA), and 40 ng/ml of dexamethasone (Sigma-Aldrich, USA). Hep-EVs were used to treat AML12 cells for 48 h at a protein concentration of 10 μg/ml. The RO-3306 compound was dissolved in dimethyl sulfoxide (DMSO) (D8371, Solarbio, China) and added to Hep-EVs at a concentration of 5 μM for a duration of 24 h then washed prior to Hep-EVs treatment.

For the wound closure assay, hepatocytes were seeded into 6-well plates at 5×10^5^ cells/well. When the cells reached 90% confluence, sterile 200 μl pipette tips were used to create cell wounds on the plates. After washing with PBS, the medium was changed into culture medium without FBS and added with 10 μg/ml of Hep-EVs dissolved in PBS to specific wells for a duration of 48 h. Cells were photographed at 0 h, 6 h, and 12 h with an inverted microscope (Leica, Germany). Quantification of the wound area was performed using the ImageJ software (NIH, USA). For the EdU labeling assay, hepatocytes were seeded into 6-well plates at 5×10^5^ cells/well. DNA synthesis was examined by 2 h EdU labeling using a commercial kFluor488 Click-iT EdU kit (KGA331, KeyGEN, China). Quantification of the positively labeled cell percentages was performed using the ImageJ software (NIH, USA).

*Integrated analysis of associations between genes and proteins*

The genes/proteins and DEGs/DEPs detected in the scRNA-seq and proteomic analyses were counted. Correlation analysis was performed by R (version 3.5.1). A nine-quadrant map was drawn based on changes in the gene expression in the hepatocyte transcriptome and the Hep-EV proteome. Quantitative and enrichment analysis of genes in each quadrant was performed.

*TEM analysis*

TEM was applied to confirm the presence of EVs in livers. All post-perfusion liver from the PHx and Sham mice were cut into pieces of about 1 mm × 1 mm × 1 mm, quickly fixed in 3% glutaraldehyde, and postfixed in a 1% OsO_4_ solution at 4°C. After fixation, the samples were dehydrated with gradient acetone and embedded in araldite (EM TP, Leica, Germany). The sliced sections applied by an ultramicrotome (EM UC7, Leica, Germany) were stained with uranyl acetate and lead citrate and were examined with 120 KV TEM (JEM-1400FLASH, JEOL, Japan).

The morphology of LT-EVs was also determined by TEM. A total of 4 μL of the EV solution, with a protein concentration of 1 mg/ml, was deposited onto a carbon-coated 400-square mesh copper grid. Ten minutes after the sample was deposited, the grid was rinsed with 10 drops of deionized water. A drop of 1% phosphotungstic acid (12501-23-4, RHAWN, China) was added to the grid to conduct the negative staining. The grid was subsequently dried naturally and visualized using the 120 KV FEI TEM (TECNAI Spirit, FEI, USA). TEM-EDS was conducted using a field emission TEM (JEOL, Japan).

*NTA experiments*

The concentration and size distribution of LT-EVs and sorted Hep-EVs were analyzed by the Nanosight NS300 system (Malvern Panalytical, UK). Data were analyzed by Nanosight NTA 2.3 Analytical Software with the detection threshold optimized for each sample and screen gain at 10 to track as many particles as possible with a minimal background. A blank 0.2 μm-filtered PBS was also run as a negative control.

*Western blot analysis*

LT-EV and Hep-EV proteins were extracted using the RIPA lysis buffer (P0013B, Beyotime, China). The BCA protein assay kit (PA115, TIANGEN, China) was used to detect the protein concentration of each sample. All samples were prepared at a final concentration of 1 μg/μl in a loading buffer (CW0027S, CwBio, China). Protein samples of 20 μg were loaded into a 4-20% SDS-polyacrylamide gel (LK206, Epizyme, USA) in the Bio-Rad Electrophoresis System to separate the proteins with different molecular weights. The proteins in the gel were then transferred to polyvinylidene difluoride (PVDF) membranes. After blocking in 5% bovine serum albumin (BSA) solution (218072801, MP Biomedical, USA), the membranes were incubated with CD63 (SC5275, Santa Cruz Biotechnology, USA), TSG101 (ab125011, Abcam, UK), Flotilin-1 (18634S, Cell Signaling Technology, USA), Golgin84 (NBP1-83352, Novus Biologicals, USA), Dnmt1 (5032T, Cell Signaling Technology, USA), Ncapg2 (24563-1-AP, Proteintech, China), Mcm3 (A1060, Abclone, China), Prg4 (PA3-118, Invitrogen, USA), or Cdk1 (ab18, Abcam, UK) primary antibodies at 4°C overnight. After incubation with the corresponding secondary antibodies (Jackson ImmunoResearch, USA) for 1 h at room temperature, the PVDF membranes were imaged using Western chemiluminescent horseradish peroxidase (HRP) substrate (Millipore, USA) with an imaging system (Tanon 4600, Shanghai, China).

*Flow cytometric analysis*

Collected LT-EVs were stained for ASGPR (sc-166633, Santa Cruz Biotechnology, USA), F4/80 (ab6640, Abcam, UK), CD11b (101201, BioLegend, USA), CD31 (102407, BioLegend, USA), GFAP (38014, SAB, China), CK19 (60187-1, Proteintech, China), CD3 (100235, BioLegend, USA), or CD19 (152420, BioLegend, USA), using primary antibodies and their isotype control, PE Mouse IgG2b,κ Isotype Ctrl Antibody (400312, BioLegend, USA), APC Mouse IgG2b,κ Isotype Ctrl Antibody (981906, BioLegend, USA), or Biotin Mouse IgG2b,κ Isotype Ctrl Antibody (401203, BioLegend, USA), at 1:100 for 1 h at 4ºC, followed by fluorescence-conjugated secondary antibodies. After washing with PBS, the percentages of positively stained LT-EVs were determined with a flow cytometer (NovoCyte; ACEA Biosciences, USA) and analyzed using NovoExpress software.

*IF staining*

Fresh liver tissue samples were fixed in 4% paraformaldehyde (PFA) (Biosharp, China) at 4°C for 4 h, washed with PBS, and dehydrated with 30% sucrose for 24 h. After being embedded in an optimal cutting temperature (OCT) compound (Leica, Germany), 10 μm cryosections were prepared with a Cryastat (CM1950, Leica, Germany). Air-dried cryosections were permeabilized by 0.3% Triton X-100 (Sigma-Aldrich, USA) for 20 min at room temperature, blocked in goat serum (Boster, China) for 30 min at room temperature, and incubated with a rabbit anti-mouse Ki67 primary antibody (ab15580, Abcam, UK), a rat anti-mouse F4/80 primary antibody (ab6640, Abcam, UK), a mouse anti-mouse TNF-α primary antibody (ab1793, Abcam, UK), a rabbit anti-mouse VCAM-1 primary antibody (A19131, Abclonal, China), a rat anti-mouse Stabilin2 primary antibody (D17-3, Medical & Biological Laboratories, Japan), a mouse anti-mouse GFAP primary antibody (38014, SAB, China), or a rabbit anti-mouse α-SMA primary antibody (ab124964, Abcam, UK) overnight at a concentration of 1:100 at 4°C. After washing with PBS, sections were then stained with an Alexa Fluor 594-conjugated donkey anti-rabbit secondary antibody (R37119, Invitrogen, USA), Alexa Fluor 488-conjugated goat anti-mouse secondary antibody (A-11001, Invitrogen, USA), an Alexa Fluor 594-conjugated goat anti-mouse secondary antibody (A-11005, Invitrogen, USA), or an Alexa Fluor 488-conjugated chicken anti-rat secondary antibody (A-21470, Invitrogen, USA). The cryosections were incubated with the appropriate fluorescence-conjugated secondary antibodies at 4°C for 1 h at room temperature at a concentration of 1:200. For F-actin staining of hepatocyte borders, after washing with PBS for three times, sections were probed with phalloidin conjugated to Alexa Fluor 488 (R37110, Invitrogen, USA) according to the manufacturer’s instructions, and counterstained with 4,6-diamidino-2-phenylindole (DAPI) (ab104139, Abcam, UK). The liver tissues were imaged by CLSM (A1plus, Nikon, Japan) and analyzed using the ImageJ software (NIH, USA).

Cultured AML12 hepatocytes were washed with PBS for three times, and fixed with 4% PFA for 30 min at room temperature, followed by washing with PBS and blocking in goat serum for 30 min at room temperature. Cells were incubated with a rabbit anti-mouse Ki67 primary antibody (ab15580, Abcam, UK) with a rat anti-mouse PHH3 primary antibody (66863-1-lg, Proteintech, China) overnight at a concentration of 1:100 at 4°C. After washing with PBS, cells were then stained with an Alexa Fluor 594-conjugated donkey anti-rabbit secondary antibody (R37119, Invitrogen, USA) with an Alexa Fluor 488-conjugated goat anti-mouse secondary antibody (A-11001, Invitrogen, USA) at room temperature for 1.5 h, followed by washing with PBS and nuclei counterstaining with DAPI (ab104139, Abcam, UK). Fluorescence imaging was carried out by CLSM (A1plus, Nikon, Japan) and analyzed using the ImageJ software (NIH, USA).

*Histological staining*

Fresh liver tissue samples were fixed in 4% PFA for 24 h and washed with running water to remove excess PFA. Then, the samples underwent dehydration through graded ethanol and were embedded in paraffin and sectioned at 5 μm per slice. H&E staining was performed with a commercial staining kit (Baso Technology, China) and images were taken using the SLIDEVIEW VS200 (Olympus, Japan).

*Statistical analysis*

All data were expressed as mean ± standard deviation (SD). Statistical comparisons between data sets were conducted with an analysis of normality and variance, followed by a two-tailed unpaired Student’s *t* test for two-group comparisons and one-way ANOVA with Turkey’s post-hoc tests for multiple group comparisons using the GraphPad Prism 9.0.0 software. Survival rates were analyzed using the Log-rank test. Difference was considered statistically significant when *p* < 0.05.

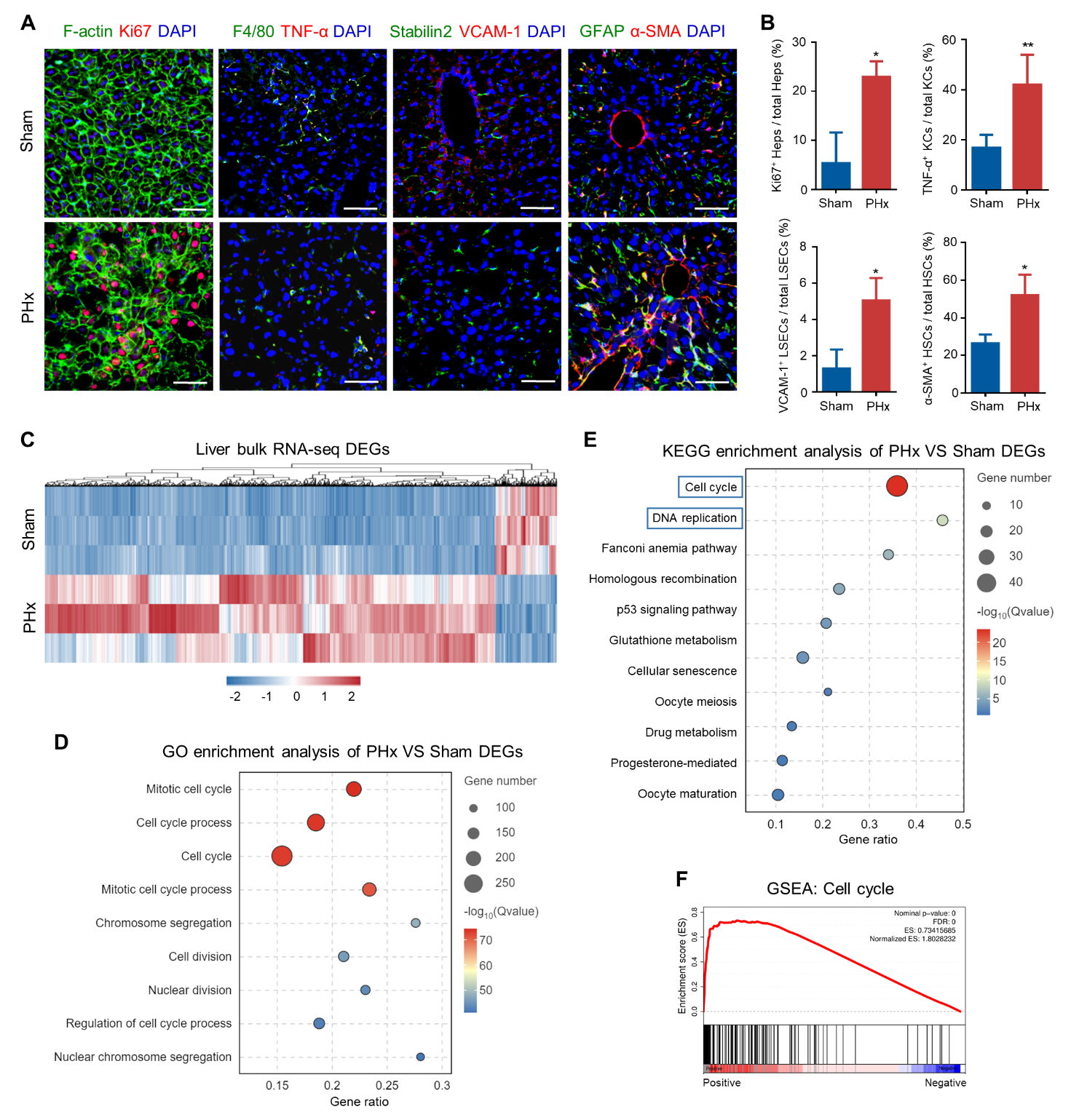

**Figure S1. Liver regeneration profiles after PHx, related to Figure 1. (A)** IF staining of hepatocyte proliferation, macrophage inflammation, LSEC and HSC activation in the liver. Bars = 50 μm. **(B)** Quantification of percentages of proliferative hepatocytes, inflammatory macrophages, activated LSECs and HSCs in the liver. Mean ± SD. n = 3 per group. *, *p* < 0.05; **, *p* < 0.01; two-tailed Student’s unpaired *t* tests. **(C)** Heatmap displaying DEGs between PHx and Sham livers. **(D)** GO enrichment analysis of DEGs in PHx over Sham livers. **(E)** KEGG enrichment analysis of DEGs in PHx over Sham livers. **(F)** GSEA analysis of DEGs between PHx and Sham livers for the KEGG term “Cell cycle”.

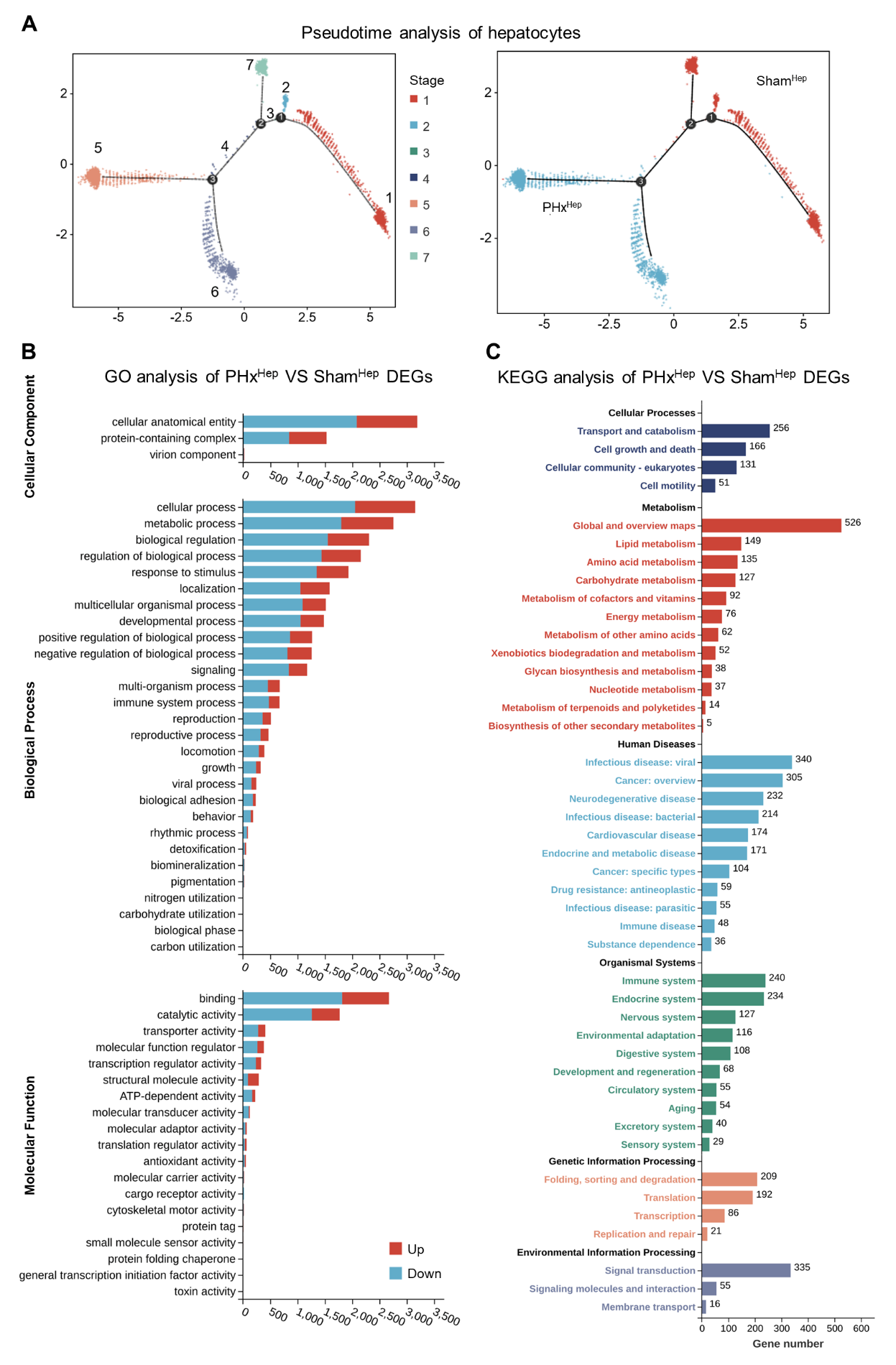

**Figure S2. scRNA-seq analysis of hepatocytes during liver regeneration, related to Figure 2.** (**A**) Pseudotime analysis plots indicating cellular trajectories of hepatocytes in PHx and Sham livers. **(B**) GO analysis of DEGs between PHx and Sham hepatocytes. **(C)** KEGG analysis of DEGs between PHx and Sham hepatocytes.

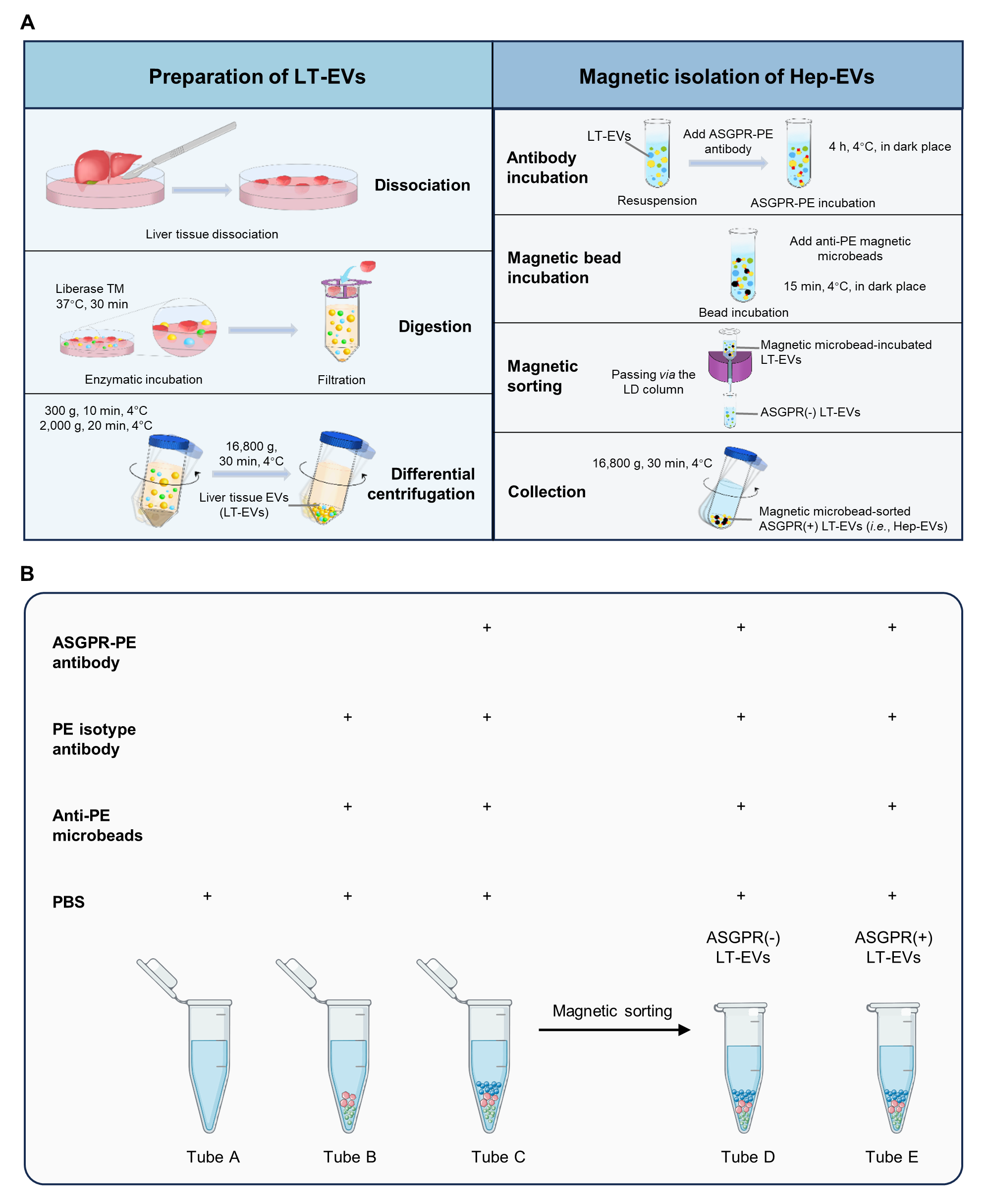

**Figure S3. Immunomagneto-activated analysis of Hep-EVs, related to Figure 3**. **(A)** Schematic overview of the protocol employing enzymatic digestion, gradient centrifugation, and immunomagnetic separation for collecting Hep-EVs from liver tissues. **(B)** Schematic diagram showing the experimental design of flow cytometric validation of Hep-EV isolation.

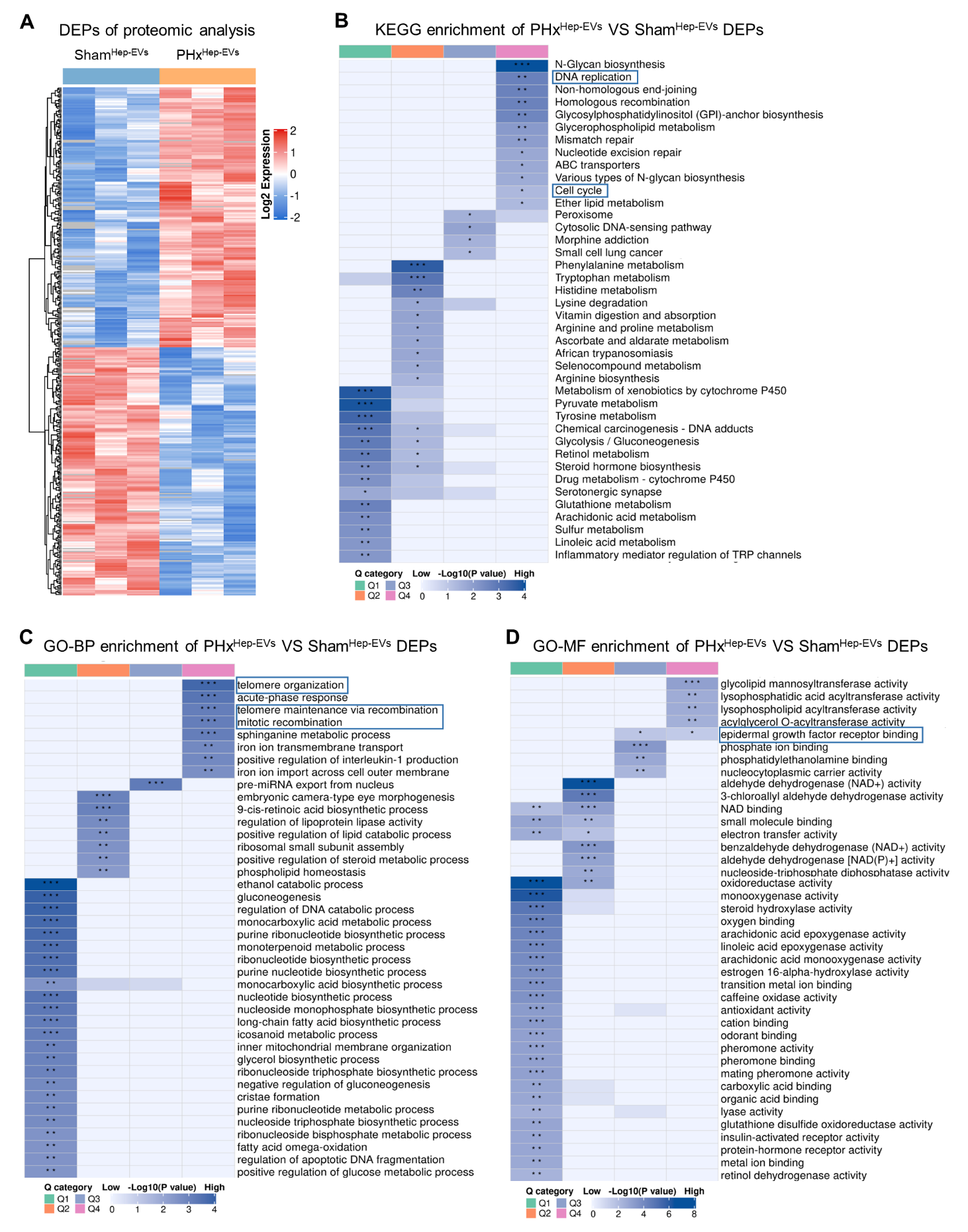

**Figure S4. Proteomic analysis of Hep-EVs during liver regeneration, related to Figure 4**. **(A)** Heatmap displaying DEPs between PHx and Sham Hep-EVs. **(B)** KEGG enrichment analysis of DEPs in PHx over Sham Hep-EVs. **(C)** GO terms in the Biological Process category of DEPs enriched in PHx over Sham Hep-EVs. **(D)** GO terms in the Molecular Function category of DEPs enriched in PHx over Sham Hep-EVs.

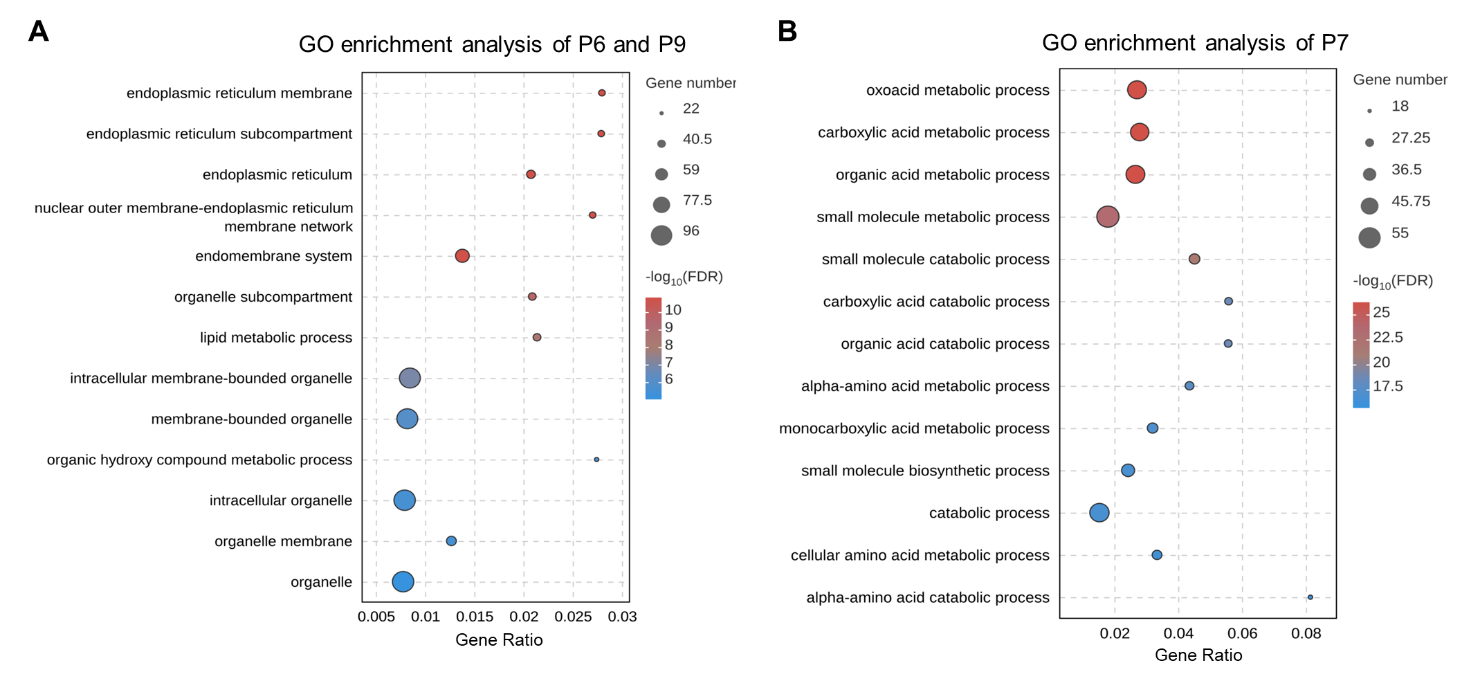

**Figure S5. Integrated analysis of hepatocyte transcriptome and Hep-EV proteome, related to Figure 5**. **(A)** GO enrichment analysis of the P6 and P9 quadrants showing specific upregulation of Hep-EV proteins rather than hepatocyte genes. **(B)** GO enrichment analysis of the P7 quadrant showing consistent downregulation of hepatocyte genes and Hep-EV proteins.

**Captions of Supplementary Tables**

**Table S1. List of DEGs identified during transcriptomic analysis**

**Table S2. List of all the proteins identified during proteomic analysis**
